# COBRA improves the quality of viral genomes assembled from metagenomes

**DOI:** 10.1101/2023.05.30.542503

**Authors:** LinXing Chen, Jillian F. Banfield

**Author notes:** To whom correspondence should be addressed: **LinXing Chen**, Address: Innovative Genomics Institute, University of California, Berkeley, CA 94720, **Jillian F. Banfield**, Address: McCone Hall, Berkeley, CA 94720.

## Abstract

Microbial and viral diversity, distribution, and ecological impacts are often studied using metagenome-assembled sequences, but genome incompleteness hampers comprehensive and accurate analyses. Here we introduce COBRA (<u>C</u>ontig <u>O</u>verlap <u>B</u>ased <u>R</u>e-<u>A</u>ssembly), a tool that resolves *de Bruijn* graph based assembly breakpoints and joins contigs. While applicable to any short-read assembled DNA sequences, we benchmarked COBRA by using a dataset of published complete viral genomes from the ocean. COBRA accurately joined contigs assembled by metaSPAdes, IDBA_UD, and MEGAHIT, outcompeting several existing binning tools and achieving significantly higher genome accuracy (96.6% vs 19.8-59.6%). We applied COBRA to viral contigs that we assembled from 231 published freshwater metagenomes and obtained 7,334 high-quality or complete species-level genomes (clusters with 95% average nucleotide identity) for viruses of bacteria (phages), ∼83% of which represent new phage species. Notably, ∼70% of the 7,334 species genomes were circular, compared to 34% before COBRA analyses. We expanded genomic sampling of ≥ 200 kbp phages (i.e., huge phages), the largest of which was curated to completion (717 kbp in length). The improved phage genomes from Rotsee Lake provided context for metatranscriptomic data and indicated *in situ* activity of huge phages, WhiB and *cysC*/*cysH* encoding phages from this site. In conclusion, COBRA improves the assembly contiguity and completeness of microbial and viral genomes and thus, the accuracy and reliability of analyses of gene content, diversity, and evolution.

## Introduction

Viruses are the most abundant biological entities on earth and play significant roles in host evolution and ecology by infecting and killing their hosts, altering host metabolisms via auxiliary metabolic genes (AMGs), and mediating horizontal gene transfer ^1–4^. The current studies of viral diversity and AMG contents are often based on genomic sequences assembled from metagenomes ^4^, most of which are partial ^5^. The diversity of viruses is extremely high ^4^, yet a relatively small fraction is represented by complete genomes ^6–8^, and only a small subset of these are for phages with genomes ≥ 200 kbp in length (which are known as jumbo phages or huge phages) ^9–15^. The lack of complete genomes often precludes the classification of extrachromosomal elements and confounds analyses of phage diversity ^6^. When complete genomes are available it is possible to evaluate phage species richness in an ecosystem, their AMG contents (thus potential geochemical impacts) ^11^, genome structure (e.g., linear vs. circular, genetic code utilization, and coding patterns) ^16^, and genome sizes ^10^.

In metagenomic analyses, a subset of *de novo* assembled contigs can be joined via their shared end sequences with a determined length ^17^, that is the max kmer (maxK hereafter) when assembled by metaSPAdes ^18^ or MEGAHIT ^19^, or maxK-1 by IDBA_UD ^20^. This is because the *de Bruijn* graph-based assemblers generally break at the positions with multiple available paths (for example, within-population variation). Thus, the fragmented contigs of a single population can sometimes be joined via manual curation to obtain complete bacterial, archaeal, and viral genomes from short-read sequenced metagenomic datasets ^9–11,17^. This process generally involves contig end extension using unplaced paired reads, so that they can be joined to other contigs from the metagenome ^17^, and manual curation is involved to evaluate the validity of joins, and eliminate chimeric joins (introduced during assembly). However, manual curation is labor-intensive and time-consuming and thus rarely included in metagenomic analysis pipelines. Binning is another strategy to better sample viral genomes from metagenomes ^21,22^, following methods used to generate bacterial and archaeal genomes from metagenomes (that is, metagenome-assembled genomes, MAGs) ^23^. Such examples of tools for binning viral fragments include vRhyme ^24^, CoCoNet ^25^, and PHAMB ^26^. However, binning algorithms are approximate and they do not improve the contiguity of individual sequences. Accordingly, we developed COBRA (<u>C</u>ontig <u>O</u>verlap <u>B</u>ased <u>R</u>e-Assembly) to detect, analyze, and join contigs from a single metagenomic assembly. COBRA evaluates coverage and paired read linkages before a join of contigs is made, following methods developed for manual curation.

We tested the ability of COBRA to improve viral genome recovery by analysis of an ocean virome dataset ^27^. This prior study combined single-molecule Nanopore long and Illumina short-read sequencing to generate 1,864 circularized viral genomes. We found that COBRA accurately joins contigs assembled from short Illumina reads alone, to generate large genome fragments and sometimes circular genomes (some were not obtained in the original study). Compared with the performance of the evaluated binning tools, almost all of the genomes generated by COBRA were accurate and not confounded by the contamination that is introduced by binning. We subsequently used COBRA to recover high-quality and circular phage genomes from 231 published freshwater metagenomes, expanding genomic sampling of phage diversity, including the diversity of huge phages, whiB-encoding actinophages, and *cysC*/*cysH* encoding phages. We showed that COBRA also improves the contiguity of whole metagenomes and draft microbial genomes from three metagenomics samples. Thus, we demonstrate that COBRA can improve and accelerate microbial and viral research.

## Results

### Simulations demonstrate the basis for joining contigs after assembly

We used simulations to investigate why and how fragmentation occurs when short read datasets are assembled using the most commonly used *de Bruijn* graphs based assemblers, i.e., metaSPAdes, IDBA_UD, and MEGAHIT (Supplementary Table 1). Specifically, our simulated data included (1) repeats within a genome (Supplementary Figs. 1 and 2), (2) regions shared by different genomes (Supplementary Fig. 3), and (3) within-population sequence variation (Supplementary Figs. 4 and 5), taking into account a range of relative abundances for cases in (2) and (3). The simulated paired-end read datasets were *de novo* assembled individually (see Supplementary Information). In the vast majority of cases where fragmentation occurred due to repeats (i.e., (1) or (2)) and all cases where it occurred due to sequence variation (i.e., (3)), the assemblers introduced end sequences (maxK for metaSPAdes and MEGAHIT assembly, or maxK-1 for IDBA_UD assembly) that could be used to suggest contig joins. We acknowledge that these joins may not be legitimate, but rather serve as hypotheses that can be evaluated using information discussed below. These findings served as the foundational knowledge that informed the development of COBRA.

### COBRA joins contigs using end overlap, constrained by sequencing coverage, and paired reads information

The thesis upon which COBRA is based is that it is possible to make joins that the assembler chose not to make so long as there is sufficient information to do so. Ideally, the assembler will not make a join that is non-unique, but some non-unique options could arise due to a single read (e.g., due to sequencing error), or a few reads (e.g., from a strain variant) that do not represent a unique part of the genome (i.e., the coverage is much lower). Thus, the first criteria that COBRA uses to evaluate potential joins based on contig end overlap is coverage. COBRA detects contigs with shared end overlap sequences of the expected length (maxK for metaSPAdes and MEGAHIT, maxK-1 for IDBA_UD), and checks if the contigs have similar sequencing coverage and are spanned by paired reads (Fig. 1, Supplementary Fig. 6). These joins are considered legitimate and the contigs are joined.

**Fig. 1.**
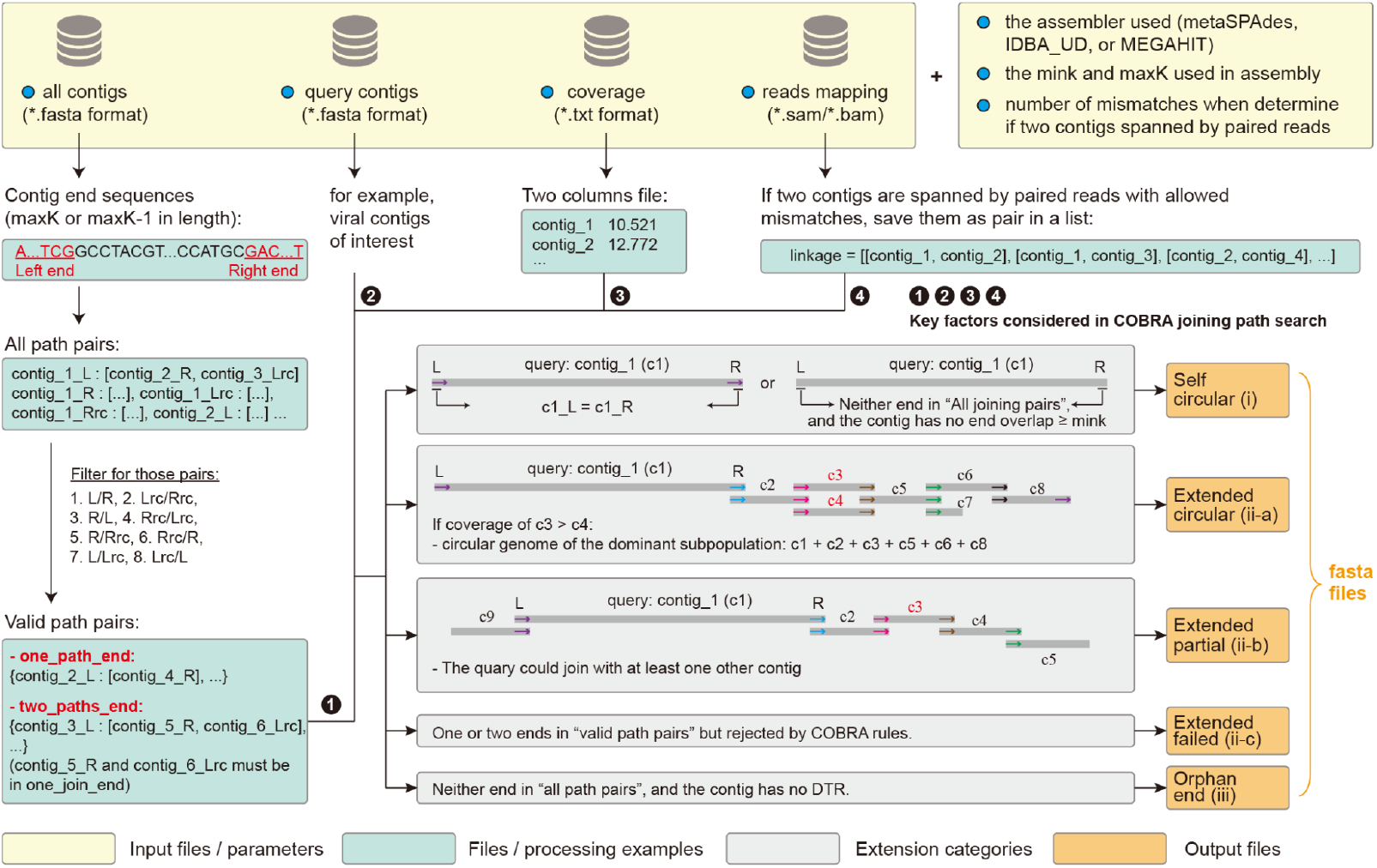
The input files and parameters, processing steps, and output files of COBRA. COBRA requires four input files: a fasta file containing all contigs from the assembly, another fasta file containing the query contigs, a two-column file with the sequencing coverage of each contig, and a read mapping file of all contigs. The parameters “assembler” and “maxk” determine the length of contig end sequences, while “mink” evaluates the end overlap sequence length of a given query to determine if it is a “self_circular” contig. See Supplementary Fig. 6 for more detailed information. The different extension categories are depicted in gray background boxes, indicating their corresponding output categories (i, ii, or iii) and associated files. For each analysis, COBRA generates five fasta files, accompanied by summary files.

In the first step, COBRA processes all contigs from a given assembly and retrieves the end sequences for each contig, that are either maxK or maxK-1 in length, depending on the specific assembler and the maxK parameter employed during assembly. COBRA first identifies all contig end pairs with the sames end sequences, considering both the end sequence and its reverse complement (rc). For example, the left end of contig 1 is referred to as contig_1_L, and the reverse complement as contig_1_Lrc. These identified pairs are then filtered to retain only those that could potentially be joined. For instance, joining the left end of one contig with the right end of another is possible if contig_1_L equals contig_2_R. In this case, the right end of contig 2 can be connected with the left end of contig 1, resulting in the combined contig 2 + contig 1. Joining the left end of one contig with the left end of another (e.g., when contig_1_L equals contig_2_L) is not possible. However, if contig_1_L equals contig_2_Lrc, the contigs can be joined, resulting in the combination of rc(contig 2) + contig 1. The filtered pairs are then examined to identify valid path pairs. COBRA labels an end for which there is only one possible join as “one_path_end”. For example, end A of contig 1 shares a sequence with only one other contig end, i.e., end B of contig 2 (Supplementary Fig. 6a, case 1). End B might share its sequence with just one other end, i.e., end A. Alternatively, end B could share its sequence with two or more ends including end A (Supplementary Fig. 6a, cases 2-4). This could indicate that the region of end B occurs multiple times in the genome or is present in two or more different genomes. COBRA labels an end as “two_paths_end” if end A shares its sequence with two other ends (end B and end C), and ends B and C share the sequence exclusively with end A (Supplementary Fig. 6b). Although this is equivalent to the reverse path for the case 2 of “one_path_end”, it is considered separately because the end under consideration for extension depends on the direction that COBRA is progressing through the contig set. Our analysis reveals that in a single assembly, the ends in categories “one_path_end” and “two_paths_end” usually account for over 99% of all ends sharing sequences with other ends.

In the second step, the users can provide a fasta file of contigs (“query contigs”) they want to try to extend (or they could try to extend all contigs in an assembly). Then, COBRA will consider each contig in the provided file and extend each end sequentially. COBRA first identifies “Self circular” (category i) contigs if neither end of a given contig has a path to any other end except the other end of itself. The script reports two possible cases of “self-circular” that differ only based on the length of the end overlap: (1) the length of the overlap is maxK (for metaSPAdes and MEGAHIT) or maxK-1 (for IDBA_UD), or (2) the length of the overlap is at least the minimum kmer length used in the assembly. Next, COBRA searches for potential joining paths for each end based on valid path pairs (either “one_path_end” and “two_paths_end”). It considers the sequencing coverage ratio between the query contig and a given candidate contig to be included in the joining path (see Supplementary Fig. 6c for details), and requires that the joins are spanned by paired reads. The path search stops when (1) the end does not share its sequence with any other end, (2) the end has three or more paths, (3) the end is within “one_path_end” or “two_paths_end”, but the coverage ratio requirement is not met and/or there is no read pair spanning the join. When a query contig is extended from one end and loops back to the other end, it is classified as “extended_circular” (category ii-a). For other queries that are extended but do not result in circularization, their status is designated as “extended_partial” (category ii-b). If at least one end of a query contig matches other ends but the join is not considered valid due to the coverage ratio and/or lack of spanning paired reads, the query is labeled as “extended_failed” (category ii-c). In cases where a query contig does not share any end sequence with others, it is assigned as “orphan_end” (category iii).

In the third step, COBRA assesses all potential joining paths identified in the second step and ensures that the paths are unique before finalizing joins. An important, but rare, case involves a query that can be extended along two (or more) seemingly unique paths. For example, paths may differ due to a strain variant, “contig 1-> contig 2a -> contig 3” and “contig 1 -> contig 2b -> contig 3”, where 2a and 2b are sequence variants. In this case, the same contigs (contig 1 and 3) have more than a single possible placement so no joins are made (“extended_failed”, category ii-c).

Additionally, COBRA searches for cases where both ends of a query contig extend into sequences that are closely related to each other. These cases are identified using a BLASTn search of the first half against the second half of a possible extended sequence (Supplementary Fig. 7). If there is a region of ≥ 1000 bp that is shared between the two halves with ≥ 70% nucleotide identity, then the query contig(s) will be assigned to the “extended_failed” category (ii-c).

In the last step, the classifications of the query contigs are compiled. Sequences in the “self_circular”category are saved, and those in the “extended_circular,” and “extended_partial” categories are joined and saved.

### CORBA accurately joins contigs fragmented during *de novo* assembly

To validate the performance of COBRA, we re-analyzed a published ocean virome sample, from which polished complete viral genomes were reported using Illumina reads and single-molecule Nanopore sequences ^27^. The short Illumina reads of the ocean virome 250M sample ^27^ were assembled using metaSPAdes, IDBA_UD, and MEGAHIT, respectively. For each assembly, we searched for the polished, complete viral genomes from the original study from all three ocean depths (1864 in total) using blastn. In addition, we recovered additional putative viral sequences using VIBRANT. Overall, we recovered 2377, 2304 and 2321 contigs, respectively (Supplementary Fig. 8, Supplementary Table 2). The three contig sets were used as the query datasets for the following COBRA analyses.

COBRA was able to categorize as circular (i.e., “self_circular”) or extend 42-56% of the contigs in the three query datasets. The remaining query contigs were assigned into the “extended_failed” (ii-c; 7-14%) or “orphan_end” (iii; 30-50%) category (Supplementary Table 2). In all but one case, the queries in the “orphan_end” category had significantly lower sequencing coverage than the queries of other categories (unpaired *t*-test; Fig. 2a), likely suggesting that these contigs broke during assembly due to insufficient reads for further extension.

**Fig. 2.**
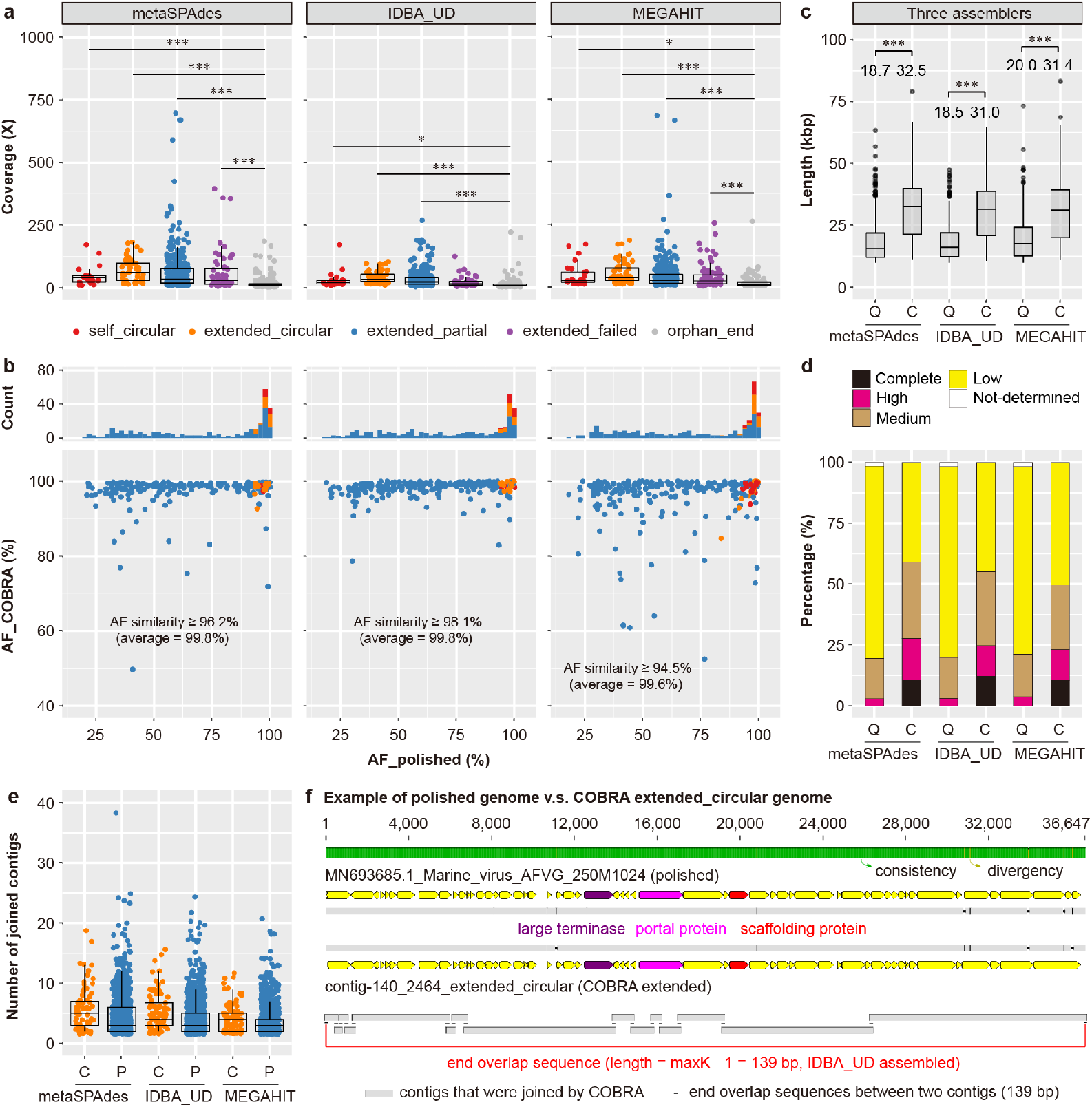
Benchmarking of COBRA using an ocean virome dataset of polished and complete viruses. (a) The coverage of query contigs in different COBRA categories. The average coverage between the “orphan_end” category and others was compared using an unpaired *t*-test (* p < 0.05, *** p < 0.001). (b) The pairwise genome alignment fraction (AF) of “self_circular”, “extended_circular”, and “extended_partial” COBRA sequences against the corresponding polished genomes (determined by BLASTn). The number of COBRA sequences in each category is plotted at the top. The minimum nucleotide similarity of the aligned region is shown, with the average similarity in brackets. See (a) for the figure legend. Comparison of the “extended_circular” and “extended_partial” sequences and the corresponding contigs joined by COBRA regarding (c) length and (d) CheckV quality. If several raw contigs were joined into one COBRA sequence, the length and quality of the COBRA sequence were counted only once. (c) The length distribution of query contigs (“Q”) and COBRA sequences (“C”). Box plots enclose the first to third quartiles of data values, with a black line at the median value. The average length of raw contigs and COBRA sequences are shown and compared using an unpaired *t*-test (*** p < 0.001). (d) The quality of query contigs (“Q”) and COBRA sequences (“C”) evaluated by checkV. (e) The number of contigs joined to generate “extended_circular” (“C”) and “extended_partial” (“P”) sequences. (f) An example of an “extended_circular” sequence compared with the corresponding polished genome. The contigs used are aligned at the bottom with their overlap shown. Box plots enclose the first to third quartiles of data values, with a black line at the median value.

We calculated the paired alignment fractions (AF) for each of the “self_circular”, “extended_circular” and “extended_partial” sequences output by COBRA and the corresponding published polished complete genomes and then we recorded the alignment fractions for each of the compared sequences (i.e., AF_COBRA and AF_polished; Fig. 2b). Generally, the higher the AF_COBRA value is the more accurate the COBRA join is considered to be. The higher the AF_polished value is, the more complete the COBRA sequence is. AF_COBRA values averaged 97.8%, 98.4%, and 97.0% for metaSPAdes, IDBA_UD, and MEGAHIT, respectively (see Supplementary Fig. 9 for examples), indicating that COBRA accurately joined the Illumina-based contigs. Lower AF_COBRA values generally occurred because (1) COBRA selected and joined strain variant contigs that represented higher-abundance subpopulations yet the corresponding polished complete genome represented lower-abundance subpopulations (Supplementary Fig. 10) or (2) COBRA sequences were very similar to, but not identical to, the corresponding polished genomes (Supplementary Fig. 11). Additionally, two MEGAHIT COBRA “extended_partial”sequences had high AF_polished (98.6% and 99.3%, respectively) but relatively low AF_COBRA (72.8% and 76.9%, respectively), as the original query contigs were longer than the corresponding polished genomes (Fig. 2b).

We assessed the length and quality of the query contigs and their COBRA sequences in the “extended_circular” and “extended_partial” categories. The average length increased by from 18.5-20.0 to 31.0-32.5 kbp (Fig. 2c), and the total number of complete/circular and high-quality genomes rose from 28-46 (3-4%) to 215-241 (23-28%) (Fig. 2d). Notably, this was achieved by joining several up to 38 contigs into a single sequence (Figs. 2e and f). These results suggested that CORBA can significantly improve the quality of query contigs. It is important to note that although COBRA only identified 31-79 “self_circular” genomes and generated 69-87 “extended_circular” genomes, this was accomplished using Illumina sequencing data, whereas the published polished complete genome reconstruction relied upon nanopore sequencing. In fact, 36-45 of the putative complete genomes generated by COBRA (i.e., “extended_circular”) were not reported in the original study.

### COBRA outperforms prevalent binning tools used for viral genome recovery

We compared the performance of COBRA to the published binning tool MetaBAT2 ^28^, which was not specifically developed for virus binning, vRhyme ^24^ and CoCoNet ^25^ that were developed for virus binning. We filtered the IDBA_UD assembly of the sample 250M ^27^ (see above) to retain contigs ≥ 2.5 kbp in length (though COBRA could work on contigs with any length), then compared them against the published polished complete virus genomes using BLASTn and used those with ≥ 99% similarity and ≥ 80% alignment coverage for binning by MetaBAT2, vRhyme, CoCoNet and contig extension via COBRA (Fig. 3a).

**Fig. 3.**
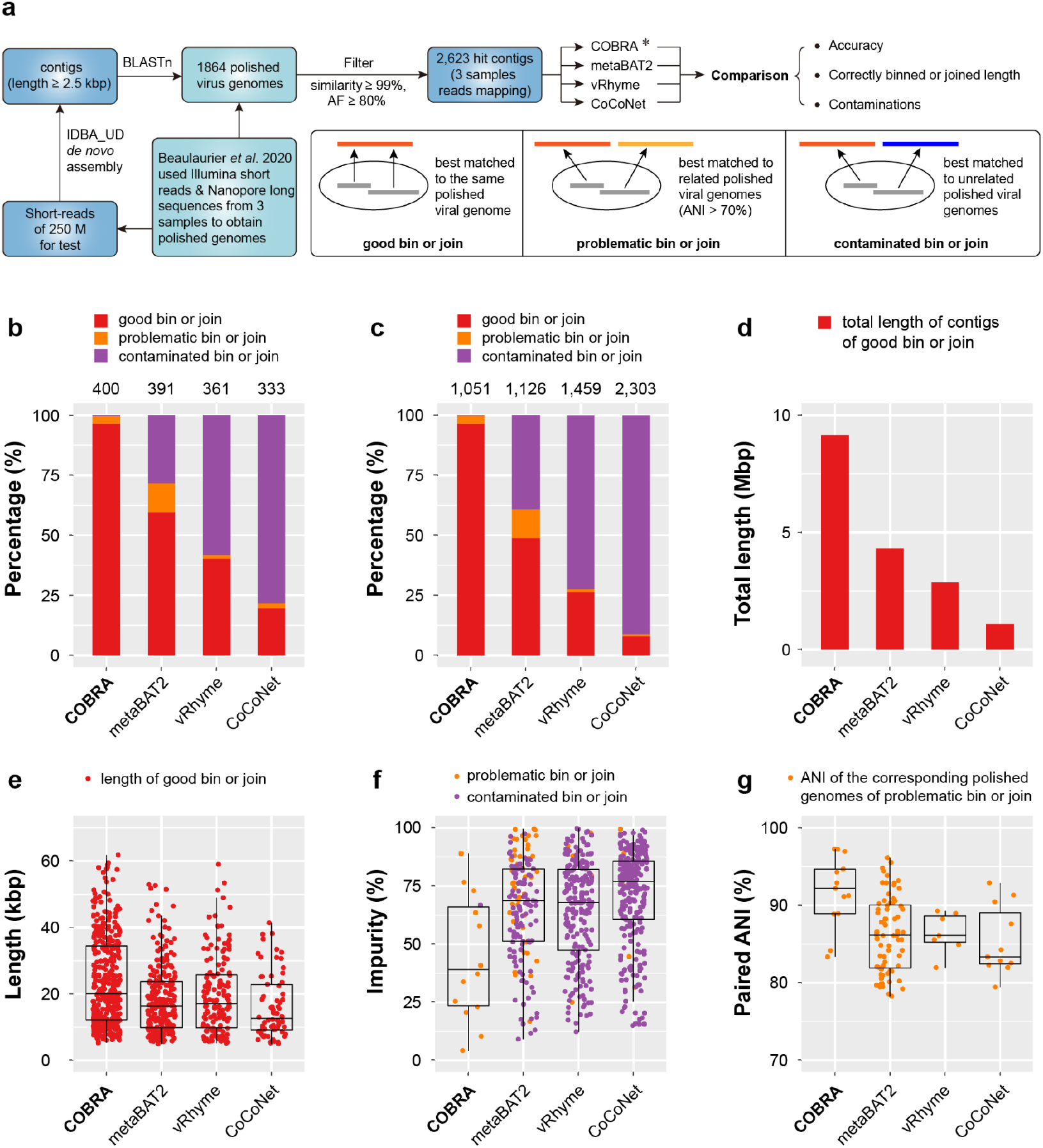
Performance comparison of COBRA and widely used binning tools. (a) The flowchart shows the comparison pipelines. The definitions of “good”, “problematic”, and “contaminated” bin or join are provided in the accompanying box. * Note that only one mapping file is needed for COBRA as input, whereas for the binning tools, the coverage profiles were obtained from all three mapping files. (b) The percentage of “good”, “problematic”, and “contaminated” bins or joins. (c) The percentage of contigs in “good”, “problematic”, and “contaminated” bins or joins. In (b) and (c), the total absolute numbers are shown at the top. For bins and joins, only those with at least two contigs binned or joined were considered and compared. (d) The total length of good bins and good joins. (e) The individual length of good bins and good joins. (f) The impurity rates of “problematic” and “contaminated” bins and joins. (g) The paired ANI of genomes that the contigs of “problematic” bins or joins were matched to. Box plots enclose the first to third quartiles of data values, with a black line at the median value.

We then compared the bins and the COBRA sequences to the published polished complete genomes to evaluate the accuracy of all approaches. To this end, we defined “good bin” and “good join”, “problematic bin” and “problematic join”, and “contaminated bin” and “contaminated join” for binning tools and COBRA, respectively (Fig. 3a, and Methods). We also compared the total length of “good bins” from binning tools to the total length of “good joins” from COBRA. In summary, as shown in Figs. 3b and c, COBRA far out-performs all binning tools in terms of its ability to recover high quality viral genomes. COBRA made 386 “good joins”, which is 1.7-5.8 times more than the contig assignments to “good bins” made by the binning tools (66-233) (Fig. 3d). The cumulative length of accurate viral sequences generated via “good joins” is 9.13 Mbp with an average length of 23.6 kbp. The cumulative length of “good bins” is 1.08-4.32 Mbp, with an average length of 16.4-19.8 kbp per bin (Figs. 3d and e).

We then investigated the problematic and contaminated bins or joins. Notably, only one out of 400 COBRA sequences was contaminated, while the binning tools generated 111-261 contaminated bins (Fig. 3f), accounting for 40-80% of all bins. In total, 13 COBRA sequences and 5-47 bins were problematic (Fig. 3f), and they were all involved closely related virus genomes with high ANI (average of 92% for COBRA, and 85-86% for bins, Fig. 3g).

### Application of COBRA to freshwater metagenomes to recover high quality viral genomes

Freshwater ecosystems contain phages that infect functionally important populations ^7,11,29^, yet their diversity is poorly understood. Here, we reanalyzed 231 published freshwater metagenomic datasets collected from 21 sampling sites around the world (Supplementary Table 3), and utilized COBRA to generate high-quality phage genomes for further analyses. We only considered sequences as queries for COBRA analysis that were at least 10 kbp in length (a total of 122,107 contigs; Supplementary Fig. 12). We filtered the COBRA output for essentially complete genomes as assessed by CheckV ^5^ and obtained 8,527 circular and 3,591 high-quality genomes for further analyses (12,118 in total; Fig. 4a). COBRA substantially improved the quality of the sequences (Fig. 4b) and the product genomes were on average 30 kbp and 25 kbp longer than the query contigs (Fig. 4c).

**Fig. 4.**
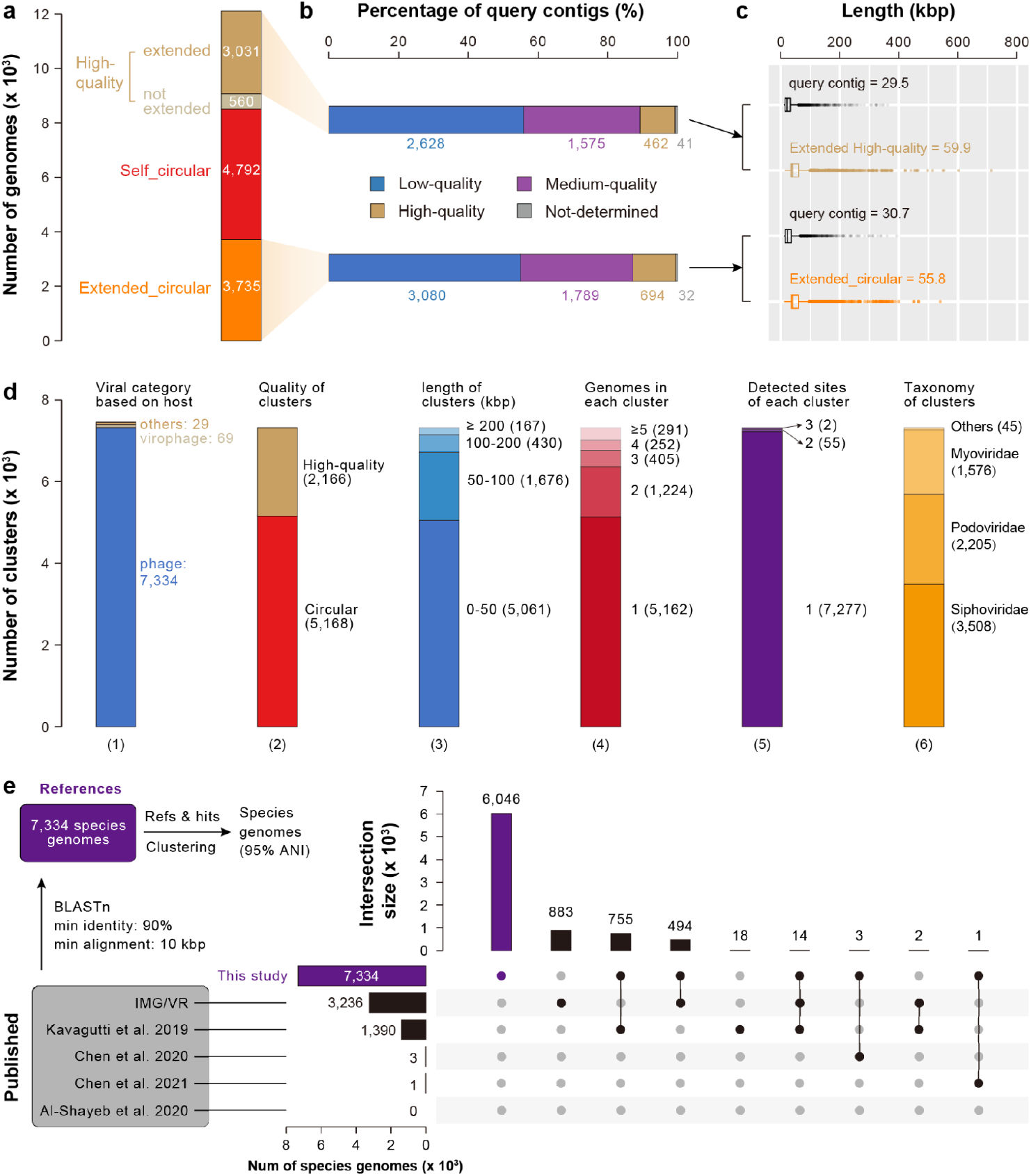
Overview of circular and high-quality phage genomes from freshwater ecosystems. (a) The number of high-quality, “self_circular”, and “extended_circular” genomes. (b) The quality of query contigs that COBRA used to generate the extended high-quality and circular genomes. The quality was evaluated by checkV. (c) The length of COBRA sequences and corresponding query contigs of “extended_partial” high-quality genomes and “extended_circular” genomes. Box plots enclose the first to third quartiles of data values, with a black line indicating the median value. (d) The clustering of viral genomes. Bar plots show (1) the number of clusters identified as phages, virophages, Eukaryotic viruses, and undetermined (“others”). The plots also show details for the 7,334 phage clusters, including (2) the number of circular and high-quality representative genomes, (3) their length distribution, (4) the number of genomes in each cluster, (5) the number of sites detected with each cluster, and (6) the taxonomic assignment of each cluster. (e) The novelty of phage species genomes identified in this study via comparison with published genomes. Of the 6,046 newly reported phage species genomes, 4,109 are circular and 1,937 are of high quality.

The 12,118 genomes were clustered into 7,432 species-level genomes (95% sequence similarity; Methods) (Fig. 4d). Of these, 69 were virophages that replicate along with giant viruses and co-infect eukaryotic cells ^30^, 29 were eukaryotes viruses or undetermined, which were excluded from further analyses. The remaining 7,334 phage species genomes included 5,169 circular and 2,165 high-quality genomes (18 of them have inverted repeats with minimum length of 50 bp), with most having a genome size of < 50 kbp (Supplementary Table 4). The majority of the phage species (70%) were represented by only one genome and 17% were represented by only two genomes. More than 99% of the phage species were detected at only one sampling site. Taxonomic analysis indicated that the phage species were mostly members of the order *Caudovirales*, with 47% belonging to *Siphoviridae*, 30% to *Podoviridae*, and 21% to *Myoviridae*. Co-analyses of the 7,334 species genomes with the previously published genomes showed that 82% of them (6,047) were novel at the species level (Fig. 4e), suggesting our analysis expanded the diversity of phages in freshwater ecosystems.

### Diversity expansion of huge phages and RNA expression of active representatives

Another motivation for developing COBRA was to obtain genomes of huge phages (or jumbo phages), which possess genomes ≥ 200 kbp in length ^31–33^. Huge phages may encode a variety of genes to exploit host replication mechanisms, and some contain CRISPR-Cas systems to assist their host in defending against smaller phages ^10^. Of the 7,334 phage species genomes identified, 167 were classified as huge phages (Fig. 3f) and their genomes underwent manual curation. In addition, 100 low- or medium-quality huge phage genomes with a minimum sequencing coverage of 20x were also chosen for manual curation. From the total of 267 huge phage species genomes, 81 were completed (error-free, gap-free, circular genomes). The largest genome was initially 712 kbp in size and ultimately reached 717 kbp after curation. To our knowledge, this is the second-largest phage genome, only 23 kbp shorter than the biggest one ever reported (735 kbp) ^10^. Two phage genomes > 800 kbp in length were reported recently ^34^ but the vast majority of these genomes are clearly bacterial sequences (Supplementary Fig. 13).

The 267 huge phages generated by COBRA have an average genome size of 285 kbp (Fig. 5a). In comparison, the original query contigs had an average size of 88 kbp, and only 102 were ≥ 200 kbp. This indicates that COBRA is highly effective in generating huge phage genomes, which are undersampled in published datasets despite their wide distribution ^10,15^.

**Fig. 5.**
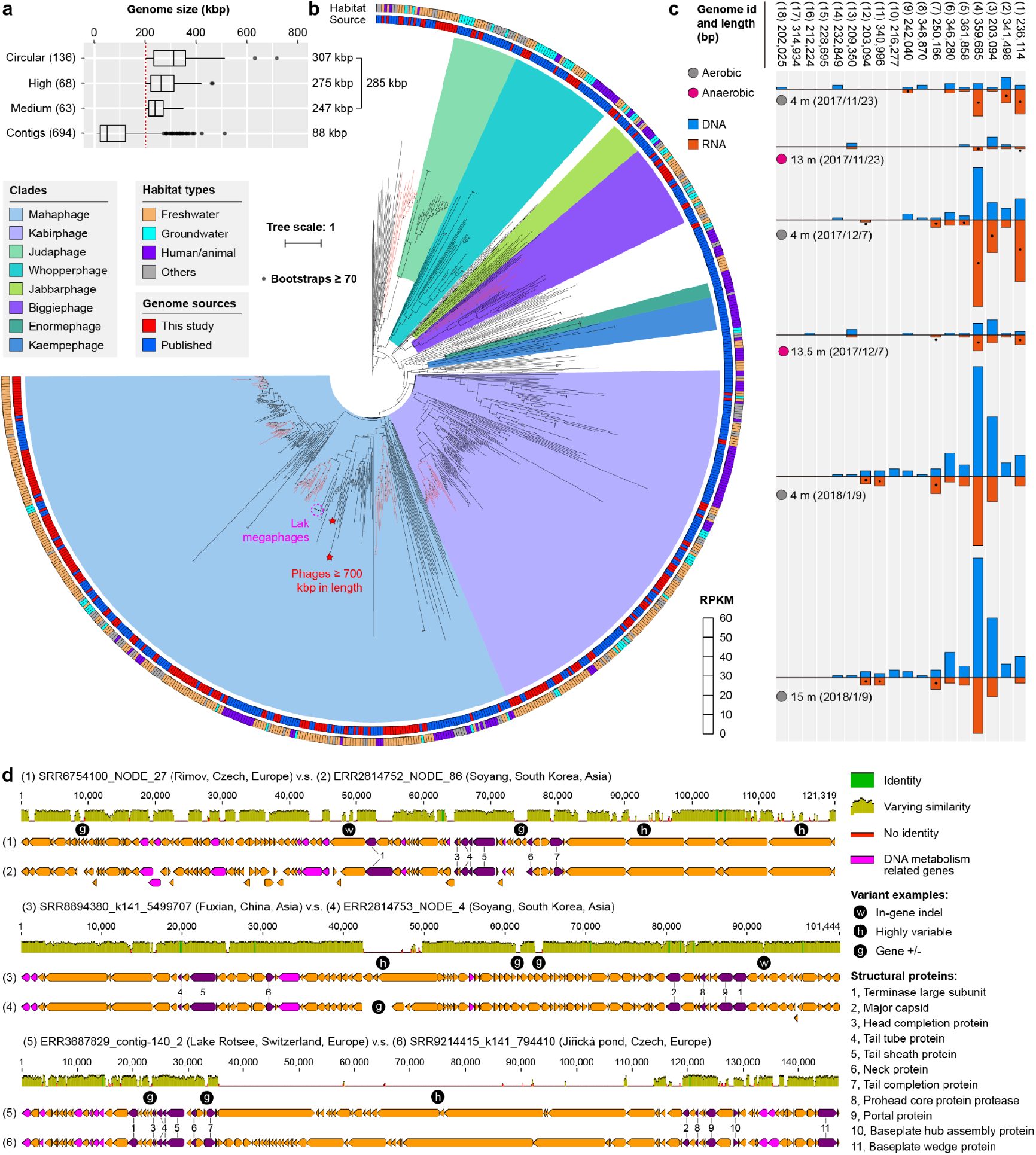
Huge phage diversity is expanded by genomes from freshwater ecosystems. (a) The number and length of huge phages newly reported in this study from freshwater metagenomes and the corresponding query contigs (≥ 10 kbp in length) joined by COBRA. Box plots enclose the first to third quartiles of data values, with a black line at the median value. (b) The phylogeny of huge phages based on the sequences of large terminase (TerL) proteins. The colored stripes in the inner ring indicate the source of genomes (published, or this study). The colored stripes in the outer ring indicate the habitats from where the phage genomes were reconstructed. The subclades with the majority (> 80%) of their genomes reconstructed in this study are highlighted in red. The two phages with genome size > 700 kbp (one published, one from this study) are indicated by red stars. (c) The detection and transcription profiles of the Rotsee Lake huge phages in the 6 samples with combined DNA and RNA analyses. The RPKM was calculated for each huge phage in each sample. A black dot indicates that the RNA RPKM is larger than the DNA RPKM of the huge phage in the corresponding sample. (d) Genomic comparison of similar huge phages from distant collecting sites. Three pairs are shown as examples (see Supplementary Fig. 14 for Mauve alignment). Structural protein genes are shown in purple, their corresponding annotations are included, and DNA metabolism-related genes are shown in pink.

To place the newly discovered huge phages within previously defined clades ^10^, phylogenetic analyses were performed based on large terminase (TerL) protein sequences (Methods). This revealed that the majority of these new huge phages are part of the Mahaphage and Kabirphage clades and are typically most similar to those identified in freshwater or groundwater habitats (Fig. 5b). Within several subclades, many genomes primarily originated from this study (highlighted in red in Fig. 5b), implying that our analyses have broadened the known diversity of huge phages.

The occurrence and transcription of the Rotsee Lake huge phages were profiled in samples collected from three-time points (Methods) (Fig. 5c). Most of the phages were detected at multiple time points, indicating their persistence in the lake. Notably, these phages were usually transcriptionally active in the aerobic water layers, with genes for structural proteins including the major capsid protein highly transcribed (Supplementary Fig. 14). Thus, huge phages appear to actively shape microbial community structure and thus biogeochemical cycles within the aerobic parts of the lake.

Comparison of huge phage genomes from different countries revealed that genes for structural proteins and DNA metabolism retain high nucleotide similarity, and that loss or gain of other genes is primarily driving their divergence (Fig. 5d, Supplementary Fig. 15).

### Expansion of the diversity of actinophages with whiB family transcriptional regulators

We assessed the impact of our phage genome collection on the sampled diversity of another group other than the huge phages. Actinobacteria are abundant in the studied ecosystems, and one of their infecting phage groups could be determined via the identification of the whiB regulators ^35^. In fact, 477 of the 7,334 new phage genomes encoded whiB (thus assigned as actinophages). The TerL based phylogeny of actinophages from the current study and published sources revealed that several subclades were defined using genomes from this study (Fig. 6a). Notably, members of certain clades appeared to encode two or more copies of whiB genes. Regardless of the source of genome data, actinophage genomes, both with and without the WhiB gene, exhibit a wide range of sizes, although only a few of them are huge phages (Fig. 6b).

**Fig. 6.**
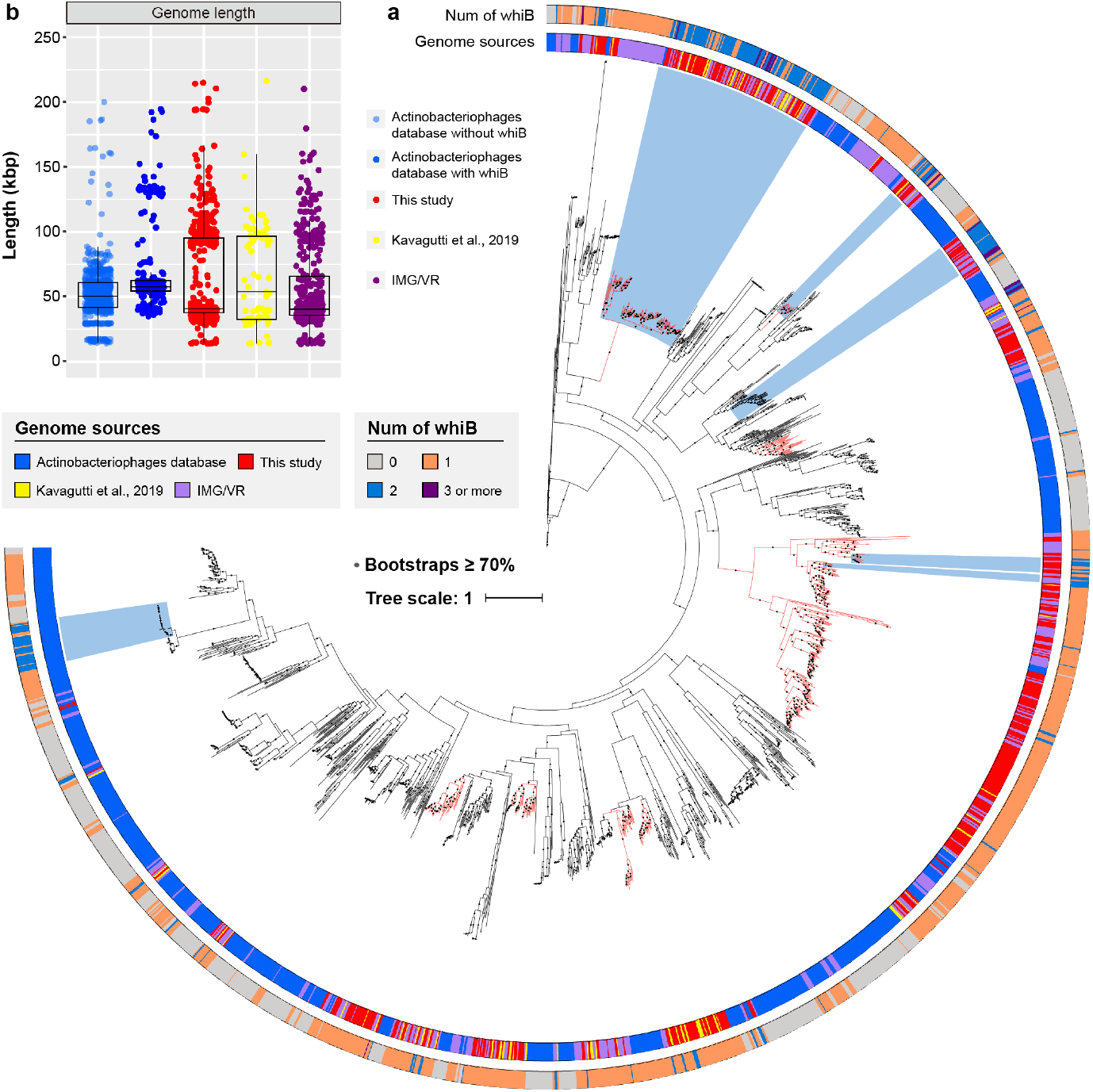
Freshwater ecosystem genomes broaden the diversity of whiB-encoding actinophages. (a) Phylogeny of actinophage genomes based on their TerL protein sequences. The numbers of whiB genes in genomes are displayed by colored stripes in the outer ring. The sources of the genomes are denoted by colored stripes in the inner ring. Subclades predominantly featuring genomes from this study are highlighted in red. Subclades, where the majority of genomes encode two or more whiB genes, are emphasized in light blue. (b) The length distribution of the genomes with or without the WhiB gene. Box plots represent the first to third quartiles of data values, with a black line indicating the median value.

The newly available high quality actinophage genomes from Lake Rotsee enabled RNA analyses of a large enough set of phages to be able to detect differences in expression patterns of different gene types. Most of the whiB genes were either inactive or transcribed at very low levels at the time of sampling. The majority of the active phages were in the late replication stage, with core structural protein-coding genes (e.g., major capsid) being expressed (Supplementary Fig. 16). In one active phage encoding two whiB genes, we observed distinct transcription levels for each gene, which likely indicates different roles for these genes in the phage’s life cycle.

### Detection of auxiliary metabolic genes in phage species genomes

We interrogated the newly available 7,334 high quality genomes to explore the inventory of Auxiliary Metabolic Genes (AMGs) in phages from freshwater ecosystems. A substantial number of AMGs were identified, the majority of which are involved in the metabolism of carbohydrates, amino acids, glycans, and cofactors/vitamins (Fig. 7a), and a few were involved in photosynthesis ^36^ and methane oxidation ^11^ were identified. Specifically, we identified 62 *cysC* genes (found in 62 genomes) and 167 *cysH* genes (found in 164 genomes) related to assimilatory sulfate reduction (Fig. 7b, Supplementary Fig. 17). These genes were generally detected in circular genomes, most of which were circularized using COBRA, and are from phages from multiple taxa (Fig. 7c). Three circular genomes each contained two *cysH* genes (refer to Fig. 7d for an example).

**Fig. 7.**
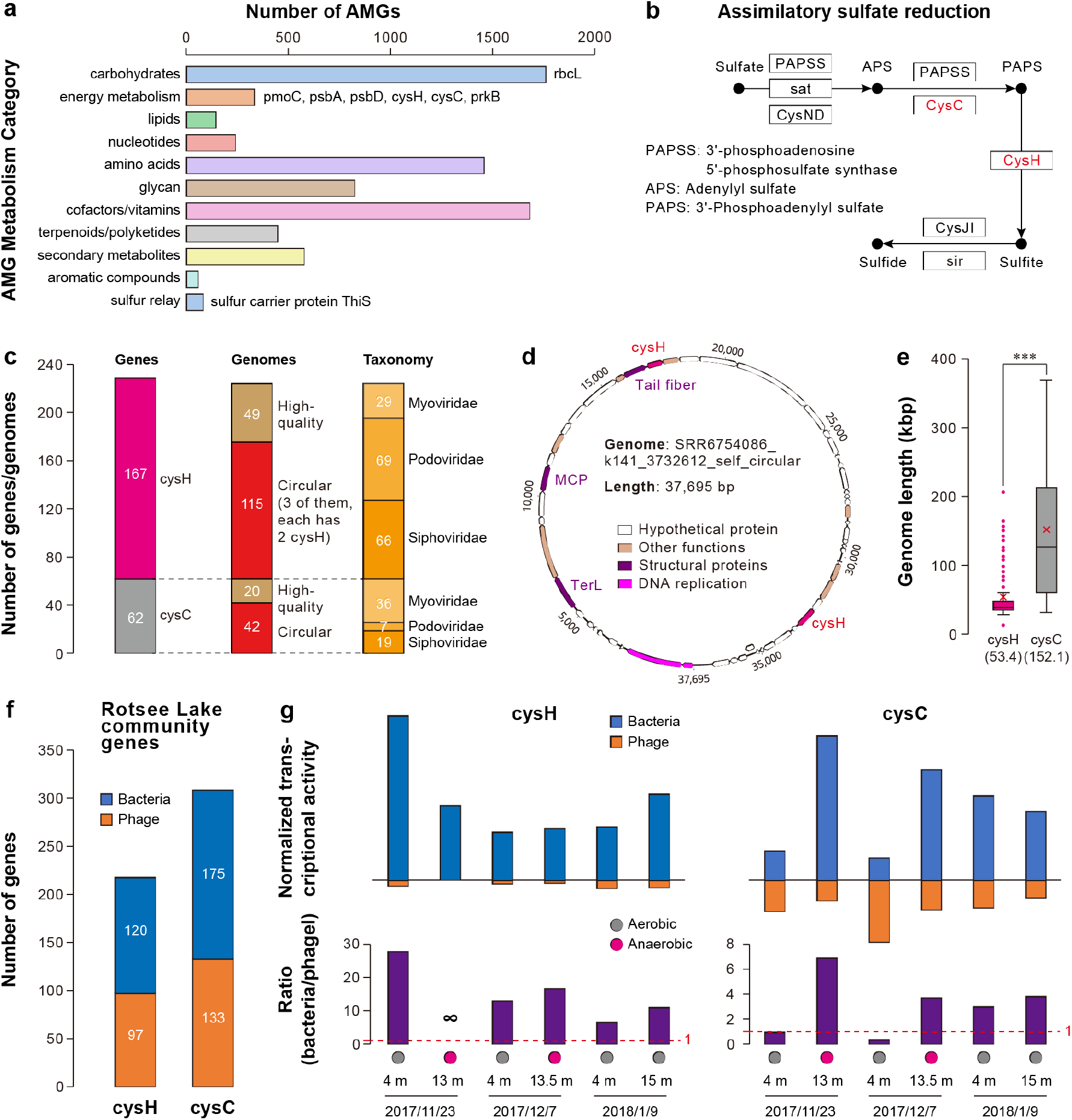
Genomic and transcriptomic analyses of phage species encoding *cysC*/*cysH* genes. (a) Summary of auxiliary metabolic genes (AMGs) identified in the phage genomes. (b) The *cysH* and *cysC* genes are involved in assimilatory sulfate reduction. (c) The quality and taxonomy of genomes encoding *cysH* and/or *cysC*. (d) One of three genomes that encode two *cysH* genes each. MCP, major capsid protein; TerL, large terminase subunit. (e) The length distribution of genomes encoding *cysH* and *cysC*. The average length of each category is indicated by a red “X”. Box plots enclose the first to third quartiles of data values, with a black line at the median value. The average length of cysH- and cysC-encoding genomes was compared using an unpaired *t*-test (*** p < 0.001). (f) The number of bacteria- and phage-encoded *cysH* and *cysC* identified in Rotsee Lake metagenomes. (g) The total normalized transcriptional activity (blue = bacterial, orange = phage) and ratio (purple) of *cysH* and *cysC* genes in Rotsee Lake metatranscriptomes. Note that the *cysC* phage transcripts are more abundant than the bacterial transcripts in the 4 m sample from 2017/11/23 and 2017/12/7.

Notably, phages encoding *cysC* genes exhibited significantly larger genome sizes (Fig. 7e). Among the 232 freshwater samples analyzed, 157 contained at least one phage with *cysC* and/or *cysH*, although the majority were only present at one of the sampling timepoints (Supplementary Table 6). Using the metatranscriptomic data from Rotsee Lake we show that under both aerobic and anaerobic conditions, the transcriptional activity of bacterial-encoded *cysH* genes generally exceeded that of phage-encoded *cysH*. However, phage-encoded *cysC* genes exhibited a greater level of transcriptional activity than their bacterial counterparts under certain aerobic conditions (Fig. 7g). These findings demonstrate that phages present in freshwater ecosystems can impact sulfur cycling via the assimilatory sulfate reduction process.

### Application of COBRA on whole metagenomes and non-viral microbial genomes

To evaluate the performance of COBRA on non-viral genomes, we applied it to three published groundwater metagenomes, each containing hundreds of MAGs from Candidate Phyla Radiation (CPR) bacteria that typically have small genomes (∼1 Mbp) ^37^, as well as from other bacteria and archaea^38,39^.

We firstly evaluated the performance of COBRA on the contigs from three metagenomic assemblies. Given the limitations of low sequencing coverage for assembly curation (Fig. 2a), we focused on the contigs with sequencing coverages of ≥ 20X and lengths of ≥ 5000 bp (fewer contigs with this length thus lower the processing time), obtaining 13,204, 7,408, and 6,827 contigs as queries for COBRA analyses respetively. COBRA substantially increased the lengths of the contigs (Fig. 8a) and significantly improved the N50, average length, and the size of the longest contig (Figs. 8b-d). Next, we evaluated COBRA’s performance on MAGs with ≤ 200 contigs (350, 310, and 179 MAGs) from the three samples). COBRA analysis significant increased the N50, average length, total length, and longest contig compared to their initial states, while simultaneously displaying a reduced number of contigs (Supplementary Fig. 18).

**Figure 8.**
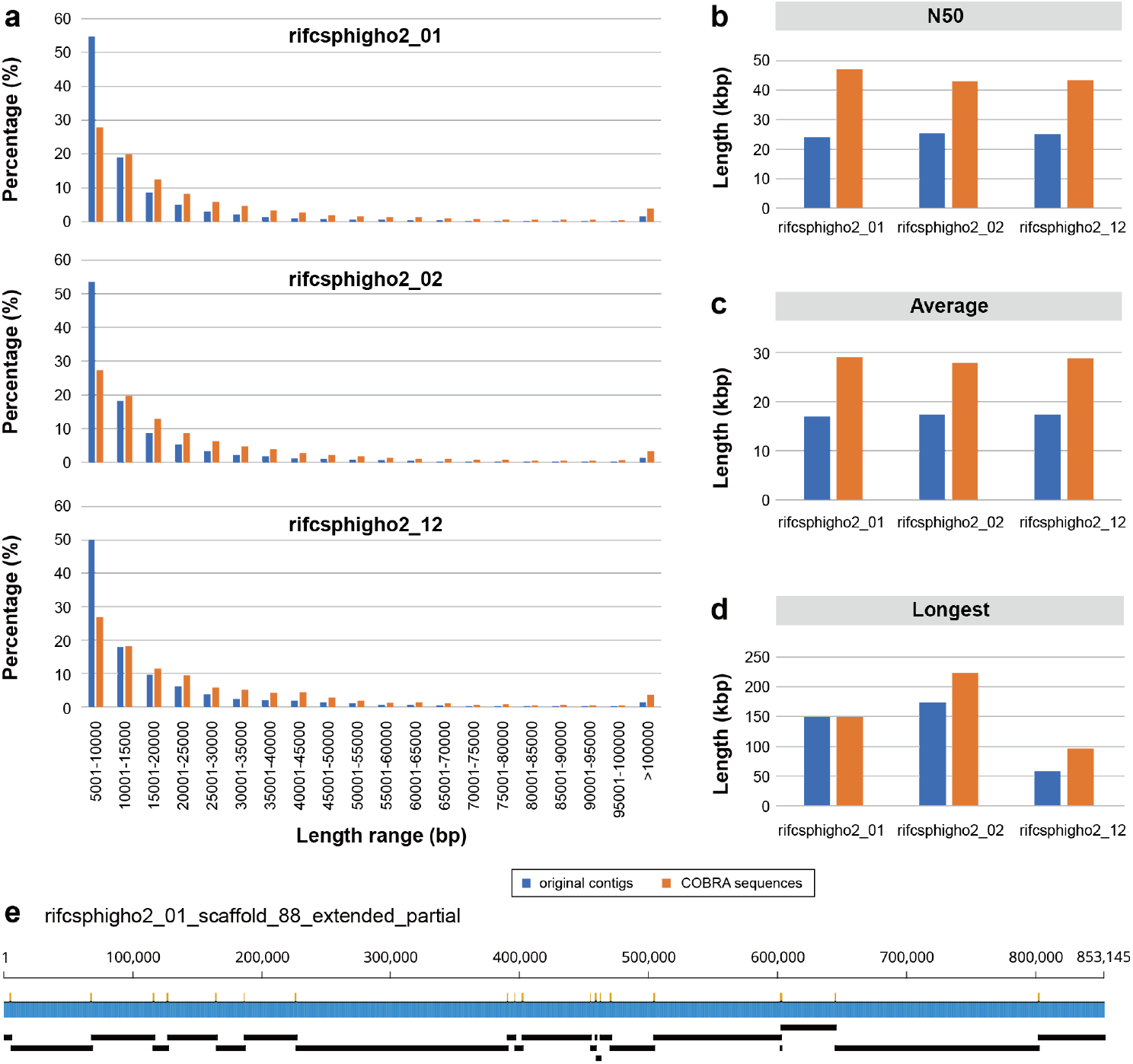
The performance of COBRA on non-viral query contigs from the metagenomic assembly. Contigs with a minimum length of 5000 bp and a minimum sequencing coverage of 20X were selected as query contigs for COBRA analyses. (a) The length distribution of original contigs (blue bars) and COBRA sequences (orange bars). (b) The N50, (c) average length, and (d) longest length of original contigs and COBRA sequences. (e) An example of where COBRA analyses joined 21 contigs into a single sequence.

## Discussion

### COBRA generates high quality viral genomes

Metagenomics is an important approach for studying viruses, the most abundant biological entities on our planet. However, fragmentation of viral genomes from metagenomes hinders understanding of their diversity and ecological significance ^5^. Fragmentation is especially problematic for viruses with large genome sizes, e.g., huge phages ^10^, and giant viruses ^13^. Previous studies primarily worked on individual viral contigs, although a few studies have attempted binning of contigs ^23^. In contrast, COBRA seeks to complete or near complete viral genomes using methods analogous to those employed in manual genome curation ^17^. COBRA should be an important tool as it can extend contigs of any length, unlike binning tools that typically require contigs of a length that is sufficient to establish reliable sequence features like tetranucleotide frequency. It can generate a single contig (sometimes, circular genomes) (Figs. 1 and 2), whereas binning tools usually obtain MAGs with two or more contigs (Fig. 3). Thus, the resulting COBRA sequences are more readily evaluated for their quality (e.g., completeness, contamination) using tools like CheckV ^5^, which does not work on bins with two or more sequences. MetaBAT2, vRhyme and CoCoNet need multiple related samples for coverage profile calculation, and PHAMB needs paired metagenome and metavirome datasets for better performance. In contrast, COBRA works efficiently on a single metagenome sample (Fig. 3). Most importantly, COBRA is much more accurate than the evaluated binning tools (Fig. 3).

An increasingly common approach to viral genome recovery involves long sequencing reads (e.g., Nanopore, PacBio). However, the methods are limited by high incidences of SNPs and INDELs ^40,41^ and long reads are unlikely to recover complete large genomes (e.g., huge phages ^10^, giant viruses ^13^. Thus, COBRA will serve as a powerful tool in viromics research.

### The expansion of phage diversity in freshwater ecosystems

Freshwater represents an ecosystem with abundant viral diversity ^6,29^ and interesting virus-encoded metabolisms ^11,42,43^. Here we demonstrate the utility of COBRA by adding over 6,000 new phage species genomes (Fig. 4). There is minimal overlap between the viral genomes reconstructed in this study and the published viral datasets (including IMG/VR; ^44^). As our study only included 231 freshwater metagenomes it is likely that viruses in the freshwater ecosystems remain underexplored. The reconstruction of huge phage genomes from distinct sampling sites allowed us to directly compare their genomes and revealed the importance of gene gain and loss in the evolution (Fig. 5). The expanded diversity of whiB-encoding actinophages suggested that the acquisition of multiple whiB genes is likely a persistent feature of several unrelated subclades (Fig. 6).

### Phage encoded *cysC*/*cysH* genes may be important in bacteria-phage interaction

The *cysC* and *cysH* genes are typically responsible for the assimilation of inorganic sulfate into organic compounds (e.g., cysteine). Some *cysC*/*cysH* encoding viruses have been reported previously (specifically, 4 genomes with *cysC* ^45^, 2 genomes with *cysH* ^46^) and very recently (4 genomes with cysH ^47^), yet their overall diversity and activity have yet to be fully understood. Here, we expand evidence for virus-driven sulfur cycling ^45,48,49^ by demonstrating a wide distribution of phages encoding *cysC*/*cysH* (Supplementary Table 6). These genes may play a role in bacterial sulfur metabolism during phage replication. Importantly, we show higher transcription of phage encoded *cysC*/*cysH* compared to bacterial *cysC*/*cysH* genes in some samples (Fig. 7).

### COBRA can also be applied to microbial genomes and whole metagenomes

Viral and microbial genomes assemblies fail for the same reasons, so COBRA can be applied generally. We demonstrate COBRA’s ability to significantly enhance the quality of generic contigs (Fig. 8) and genome bins (Supplementary Fig. 18). When applying it to large metagenomic assemblies prior to binning, we recommend using COBRA only to extend the subset of longer contigs.

## Conclusion

COBRA was developed to improve the quality of genomes assembled from short read metagenomes and its value demonstrated for viral genome recovery. Specifically, we recovered thousands of circular and high-quality new phage species genomes and hundreds of new genomes for huge phages. We anticipate that COBRA will be broadly used to extend the sequences and thus the quality of genomes assembled from metagenomes.

## Methods

### Simulated genomes for evaluation of contig breaking rules in de novo assembly

To evaluate how the assemblers of IDBA_UD, metaSPAdes, and MEGAHIT will fragment the contigs during assembly in dealing with intra-genome repeats, inter-genome shared region, and within-population variation (i.e., local variation), we simulated the artificial genomes using Geneious Prime ^50^ for different cases that are described in detail in Supplementary information. In each case, the artificial genomes were simulated for Illumina paired-end reads using InSilicoSeq with the “HiSeq” error model, which generated paired-end reads in the length of 126 bp. The simulated reads were then assembled using IDBA_UD (peng et al. 2012) (“mink = 20, maxk = 100, --step = 20, --pre_correction”), metaSPAdes version 3.15.1 ^18^ (“-k 21,33,55,77,99”), and MEGAHIT version 1.2.9 ^19^ (“--k-list 21,29,39,59,79,99”). The obtained contigs from each assembly of each case were manually checked for breaking points and the possibilities of joining via their end sequences with a determined length (i.e., 99 bp), which are shown in detail in Supplementary information.

### Evaluation of contig breaking rules in *de novo* assembly using simulated genomes

To assess how IDBA_UD, metaSPAdes, and MEGAHIT fragment contigs during assembly when confronted with intra-genome repeats, inter-genome shared regions, and within-population variation (local variation), we generated artificial genomes using Geneious Prime ^50^. Detailed descriptions of each case can be found in the Supplementary information. For each case, artificial genomes were simulated for Illumina paired-end reads of 126 bp in length using InSilicoSeq ^51^ with the “HiSeq” error model. Subsequently, the simulated reads were assembled using the following parameters: IDBA_UD ^20^: “mink = 20, maxk = 100, --step = 20, --pre_correction”; metaSPAdes ^18^ version 3.15.1 18: “-k 21,33,55,77,99”; MEGAHIT ^19^ version 1.2.9 19: “--k-list 21,29,39,59,79,99”. The resulting contigs from each assembly in each case were manually inspected for breaking points and the potential for joining via their end sequences, with a specified length of 99 bp.

Further details can be found in the Supplementary information.

### Benchmark CORBA using a previously published ocean virome dataset

Three virome datasets with samples collected from different depths (i.e., 25 m, 117 m, and 250 m) were reported previously. The authors sequenced the extracted DNA using both Illumina paired-end reads (150 bp in length) and also Nanopore single-molecule reads ^27^. With these reads, the authors detected and polished complete viral genomes, mostly from the sample collected at 250 m. This dataset was used to benchmark the performance of COBRA. To evaluate the performance of COBRA, the raw reads of the 250 m sample were downloaded from NCBI and trimmed using (https://github.com/najoshi/sickle) using default parameters to remove low-quality bases. The adapter sequence and other contaminants were detected and excluded using bbmap (https://sourceforge.net/projects/bbmap/). The trimmed reads were assembled using IDBA_UD (peng et al. 2012) (“mink = 20, maxk = 140, --step = 20, --pre_correction”), metaSPAdes version 3.15.1 ^18^ (“-k 21,33,55,77,99, 127”), and MEGAHIT version 1.2.9 ^19^ (“--k-list 21,29,39,59,79,99,119,141”). For each assembly, the contigs with a minimum length of 10 kbp were compared against the polished viral genomes reported in ^27^ using BLASTn, the hits with a minimum nucleotide similarity of 97% and minimum alignment length of 10 kbp were retained as queries for COBRA analyses. The quality reads of each sample were respectively mapped to all the contigs of the corresponding sample using Bowtie2 version 2.3.5.1 with default parameters ^52^. The sequencing coverage of the contigs was determined using the “jgi_summarize_bam_contig_depths” function from MetaBAT version 2.12.1 ^28^ and transferred to a two columns file using in-house Perl script. COBRA analyses were performed for BLASTn hits contigs from each assembler, with a mismatch of 2 for linkage of contigs spanned by paired-end reads, the maxk and assembler were flagged according to that used in assembly. The average nucleotide identity (ANI) analyses between COBRA sequences and polished genomes were performed by fastANI version 1.3 ^53^, and the alignment fraction (AF) was calculated accordingly. The quality of viral genomes was evaluated by checkV ^5^.

### Comparison the performance of COBRA and binning tools

We compared the quality of sequences joined by COBRA to bins generated by various binning tools, namely MetaBAT2 ^28^, vRhyme ^24^, and CoCoNet ^25^. Using the IDBA_UD-assembled contigs (≥ 2500 bp) from the 250 M ocean virome sample, we searched for contigs that exhibited ≥ 99% nucleotide similarity and ≥ 80% alignment coverage with polished genomes, resulting in a set of 2,632 contigs termed “query contigs”. These query contigs were extended using COBRA, and also binned using the aforementioned binning tools. For coverage calculation, the quality reads from the virome samples (25M, 117M, and 250M) were mapped to the query contigs individually using Bowtie2 version 2.3.5.1 with default parameters ^52^. The coverage profiles derived from all three mapping files were used as input for the three binning tools. However, COBRA only utilized the mapping file and coverage profile of the 250M sample. Each bin contained a minimum of two contigs, and if a bin contained only one contig, it was assigned as “unbinned.”

To evaluate the accuracy of COBRA joins and sequences represented by bins, we matched the joined contigs and bins back to the polished genomes. For the binning tools, if all the contigs from a given bin were best matched to the same polished genome, the bin was termed as a “good bin”. For COBRA, if all the contigs joined into a COBRA sequence were best matched to the same polished genome, the join was termed a “good join”. If some of the contigs matched to one polished genome (genome a), and some others best matched to another one (genome b), when genome a and genome b shared ≥ 70% ANI (determined by fastANI version 1.3 ^53^), the bin was termed as “problematic bin” (for those from binning tools), and the join as “problematic join” (for those from COBRA). To determine the extent to which the “problematic bin” or “problematic join” was affected by contigs from related (sub)populations, we also compared the ANI of the corresponding matched polished genomes.

For a given bin or join, if some contigs matched to one polished genome (genome a), and some others best matched to another one (genome b), when genome a and genome b shared < 70% ANI (determined by fastANI version 1.3 ^53^), it was termed as “contaminated bin” or “contaminated join”, respectively. To determine the contamination rate of the “contaminated bin” or “contaminated join”, we calculate the total length of the bin or join (Total_len), and also the total length of contigs best matched to each of the polished genomes, then picked up the polished genome match with the maximum total length of contigs (i.e., Max_len). We calculated the contamination rate of the contaminated bin or join as below, in which “Num_polished” is the total number of matched polished genomes. By doing so, the theoretical maximum contamination rate of a contaminated bin or join is normalized (i.e., 100%).

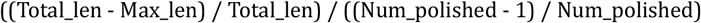

For example, where the total length of the bin or join is 100 kbp and the contigs are matched to two polished genomes, with one polished genome best matching contigs with a total length of 60 kbp and the other polished genome matching contigs has a total length of 40 kbp, Total_len = 100 kbp, Max_len = 60 kbp, Num_polished = 2, thus the contamination rate = 80%.

### Collection and analyses of published freshwater metagenomic datasets

The freshwater metagenomic datasets from two previously published studies ^7,54^ were used. The raw paired reads were downloaded from NCBI and filtered to remove any low-quality reads/bases and adaptors and other contaminants as described above. The *de novo* metagenomic assembly was first performed using the quality reads by IDBA_UD ^20^ (“mink = 20, maxk = 140, --step = 20, --pre_correction”), or metaSPAdes version 3.15.1 ^18^ (“-k 21,33,55,77,99,127”). If the RAM of our computing server was not sufficient to assemble the reads of a given sample, it was assembled using MEGAHIT version 1.2.9 ^19^ (“--k-list 21,29,39,59,79,99,119,141”). If a given sample could not be assembled using any of the three assemblers, it was excluded from the analyses. The assembler details for each dataset are shown in Supplementary Table 3.

The generated contigs with a minimum length of 10 kbp from each assembly were predicted for viral sequences using VIBRANT ^55^ using default parameters. The identified lysogenic and lytic virus contigs by VIBRANT were used as queries for COBRA analyses. A max mismatch of 2 in each read was set to identify the linkage of contigs spanned by paired-end reads, and the minK, maxk and assembler were flagged according to what was used in assembly (Supplementary Table 3).

### Filtering of COBRA sequences and evaluation of assembly gaps

The COBRA sequences from all 231 freshwater metagenomic datasets were evaluated by checkV (version 0.7.0) ^5^. The “self_circular” and “extended_circular” COBRA genomes and those identified as “High-quality” by checkV were retained for further analyses. To evaluate and fix the assembly gaps, we checked the genomes by parsing the reads mapped to them (with Bowtie2 as described above) using a custom script named “gap.check.py” (TBA). The script filtered the mapped reads to allow 2 mismatches for each read, for a region in a given genome sequence without any base mapped, the region was replaced by 10x Ns. The resulting sequences were used for further analyses.

### Genome completeness evaluation of the query contigs

To determine the extent to which COBRA raised the quality of the viral genomes we evaluated the original query contigs that were joined into “extended_partial” or “extended_circular” genomes, using checkV (version 0.7.0) ^5^. The percentages of original contigs assigned by checkV to “Low-quality”, “Medium-quality”, “High-quality”, and “Not-determined” were profiled and shown in Fig. 4b.

### Clustering of viral sequences

The quality viral sequences were clustered at the species level using the rapid genome clustering approach provided by checkV ^5^ (available at https://bitbucket.org/berkeleylab/checkv/src/master/). The clustering parameters were set as follows: -perc_identity = 90 (for BLASTn), --min_ani = 95, --min_qcov = 10, and --min_tcov = 80 (for aniclust.py). The quality viral sequences were clustered into species level clusters. Among these representative sequences, 6,430 had no assembly gaps, 815 had one gap, 195 had two gaps, and 71 had three or more gaps. It is worth mentioning that any identified gaps were filled with 10x Ns during the clustering process.

### Identification of Eukaryotes viruses, virophages, and phages

The protein-coding genes were predicted using Prodigal (-p meta) ^56^. The Eukaryotes viruses were identified by searching the core structural protein sequences via BLASTp against the RefSeq database ^57^. The virophage sequences were identified by searching their major capsid proteins (MCPs) against the virophage-specific HMM databases reported previously ^30^ using hmmsearch ^58^ version HMMER 3.3 (-E = 1e-6). Those sequences with virophage-specific MCP hits were confirmed by building a tree with the MCPs from reference virophage sequences published previously ^30,59^.

### Identification of new species phage genomes obtained in this study

The viral genomes from several published datasets were included for comparison, including the IMG/VR ^44^, the huge phage across ecosystems ^10^, and the complete viral genomes from freshwater metagenomes ^7^, the pmoC-phages ^11^, and bS21-phages ^60^, these genomes were termed as “viral_refs”. The “viral_refs” genomes were first searched against our cluster representative genomes using BLASTn with a minimum e-value of 1e-50 and a minimum similarity of 90%. The BLASTn results were parsed to retain those with at least one hit with a minimum alignment length of 5,000 bp, and the corresponding genomes were extracted for genome clustering. If a given phage genome from the clusters could clustered with any of the “viral_refs”, it was labeled as “reported”, otherwise as “new species genome”.

### Huge phage analyses

The subset of representative phage genomes with a minimum length of 200 kbp were classified as huge phages. To include more huge phage genomes from the freshwater datasets, we checked the low-quality and medium-quality genomes for huge phages, and manually curated some of them. Protein-coding genes were predicted from them using Prodigal version 2.6.3 (-m -p meta) ^56^. The predicted proteins were searched using BLASTp (e-value threshold = 1e-5) against the TerL proteins from the huge phages reported previously ^10,14^. The BLASTp hits were confirmed using the online HMM search ^61^. The phylogenetic tree was built including all the confirmed TerL hits and the above mentioned previously published huge phage TerL using IQ-TREE version 1.6.12 (-bb = 1000, -m = LG+G4) ^62^.

To evaluate the abundance of each huge phage in each of the samples from Lake Rotsee (Fig. 5c), RPKM (reads per kilobase per million reads mapped) was calculated as follows, RPKM = N_phage_ / (L_phage_/1000) / (N_sample_/1000000), where N_phage_ is the number of reads to the phage genome, L_phage_ is the length of the phage genome (bp), and N_sample_ is the number of reads mapped to the whole metagenome-assembled contig set. The DNA read mapping to genomes or contigs was performed by Bowtie2 (version 2.3.5.1) ^52^ with default parameters excepting -X = 2000, and filtered using the pysam Python module ^63^ to allow 0 or 1 mismatch for each mapped read. RPKM calculation of RNA reads to phage genomes was performed in the same way.

### Analyses of actinophages

We searched the phages infecting Actinobacteria (i.e., actinophages), which are abundant in freshwater ecosystems, by searching for the whiB gene ^35^ via blastp search against NCBI RefSeq whiB protein sequences and by manual validation using the online HMM search tool (www.ebi.ac.uk/Tools/hmmer/search/). We deteremined the subset of the recovered genomes encoded whiB that have been reported previously ^7^. A total of 4,288 (519 high-quality species genomes) from IMG/VR and 158 (79 species genomes) from Kavagutti et al. ^7^ were included in our analyses, along with 4,070 actinophage genomes (1,116 encode whiB) from “The Actinobacteriophage database” (https://phagesdb.org/). The entire set were clustered to identify distinct species genomes as described above (see section “Clustering of quality viral sequences”). The TerL protein sequences from all the sources were used to reconstruct a phylogenetic tree to show the expansion of whiB-encoding actinophages in this study. The TerL sequences were aligned using MAFFT v7.453 ^64^ with default parameters, and trimAL ^65^ was used to filter the alignment to remove those columns containing more than 80% of gaps, the tree was built using IQ-TREE version 1.6.12 ^62^ with 1,000 bootstraps and the “LG+G4” model.

### Transcriptional activity analyses

For the analysis of viral metabolic gene expression in situ, RNA reads obtained from Rotsee Lake samples were utilized. Metagenome-assembled contigs with a minimum length of 5 kbp were examined using checkV ^5^ and VIBRANT ^55^ to identify bacterial and viral *cysC* and *cysH* genes. Only contigs with consistent predictions (either bacteria or virus) from both checkV and VIBRANT were retained. The RNA reads from each sample were mapped to the corresponding contigs harboring *cysC* and/or *cysH* genes. Subsequently, the transcriptional activity of each gene was normalized and summed separately for those encoded by bacteria and phages. The ratio of total transcriptional activity between bacteria and viruses was calculated individually for *cysC* and *cysH* in each sample.

### COBRA analyses of whole metagenomes and microbial genomes

To assess the effectiveness of COBRA on whole metagenomes and microbial genomes (binned MAGs), we chose three groundwater metagenomic assemblies as representatives. In the case of whole metagenomes, we applied filters to retain contigs with a minimum sequencing coverage of 20X and a minimum length of 5000 bp. This length threshold was selected to expedite COBRA analyses due to the large number of contigs in these three assemblies. Consequently, we obtained 13,204, 7,408, and 6,827 contigs as COBRA queries for the respective metagenomic samples. These retained contigs were then utilized as queries (-q) for COBRA analyses. Subsequently, we calculated and compared the length distribution, N50 length, average length, and largest length (longest contig) of sequences before and after COBRA analyses. For microbial genomes, we selected bacterial and archaeal MAGs with a maximum of 200 contigs from each of the three metagenomes for further examination. The contigs from the corresponding MAGs were combined into a single file as queries (-q) for COBRA analyses. We subsequently calculated and compared the N50 length, largest length, average length, total length, and the number of contigs for each MAG before and after COBRA analyses.

## Data availability

The high-quality and complete genomes obtained from the 231 freshwater metagenomes are available at figshare via https://figshare.com/articles/dataset/viral_genomes_fasta/23282789. The data supporting the findings of this study are available within the paper and its supplementary information files.

## Code availability

COBRA is available as an open-source Python program on GitHub (https://github.com/linxingchen/cobra.github.io).

## Acknowledgements

The study was supported by NSERC Canada and Syncrude Canada (grant no. CRDPJ 403361-10), the Chan Zuckerberg Biohub and the Innovative Genomics Institute at the University of California, Berkeley, and the Genomic Science Program (GSP) LLNL ‘Microbes Persist’ Soil Microbiome Scientific Focus Area SCW1632 from the U.S. Department of Energy (DOE), Office of Biological and Environmental Research.

## Author contributions

L.X.C. and J.F.B. conceived and designed the study. L.X.C. developed the tool, wrote the script, and conducted simulated analyses, metagenomic assembly, viral contig identification, compared tools and performed phylogenetic, genomic and RNA expression analyses. L.X.C. drafted the manuscript, and both authors contributed to manuscript revisions.

## Competing interests

J.F.B. is a co-founder of Metagenomi.

## Notes

https://github.com/linxingchen/cobra.github.io

### Competing Interest Statement

The authors have declared no competing interest.

### Summary of Updates

Minor modifications in the Abstract section, and with updated data availability.

