## Supplementary Figures for "COBRA improves the quality of viral genomes assembled from metagenomes" (from another version of this preprint: version 1, 2023-06-01)

#### Supplementary Information
#### for

### COBRA improves the quality of viral genomes assembled from metagenomes

##### The simulation of genomes with repeats and variations.

To test how metaSPAdes, IDBA_UD, and MEGAHIT assemble short reads from genomes with **intra-genome repeat regions** (including direct terminal repeats, DTRs), **inter-genome shared regions**, and **within-population variation** (Supplementary Table 1), we obtained simulated genomes for each case and generated in-silicon Illumina HiSeq paired-end reads (126 bp X 2) using InSilicoSeq (Gourlé et al. 2019). The reads were *de novo* assembled using the three assemblers individually. The assembled results were compared against the simulated genomes, and the number of contigs from the assemblies, and the end sequences between contigs were analyzed for overlaps. See below for details of each simulated case.

**Intra-genome repeats.** We obtained simulated genomes with a region that repeats twice and three times, at both distant (means the repeat regions are distant from each other) and consecutive circumstances (means the repeat regions are next to each other) (Supplementary Fig. 1, Supplementary Table 1).

If the repeat length is shorter than the maxK used in assembly (Supplementary Fig. 1, Supplementary Table 1), only two assemblies generated fragmented contigs, (1) three times repeats that are distant from each other in the simulated genome, and assembled by MEGAHIT, and (2) three times of repeats that are consecutive to each other in the simulated genome, and assembled by IDBA_UD. For all the other cases, only one contig was generated in each, however, MEGAHIT sometimes generated only one copy of the consecutive repeats.

If the repeat length is longer than the maxK used in assembly (Supplementary Fig. 2, Supplementary Table 1), the assemblers usually generated fragmented contigs excepting, (1) two times repeats that are distant from each other in the simulated genome, and assembled by metaSPAdes, (2) two times of repeats that are consecutive to each other, and assembled by all three assemblers, while only metaSPAdes got the correct copy number of the repeats, and (3) three times of repeats that are consecutive to each other, and assembled by MEGAHIT, while with only one copy of the repeat in the contig. In all the other assemblies, two or three contigs were generated, and the length of the contig end sequences between contigs is expected (i.e., maxK or maxK-1).

We also obtained simulated genomes with DTR that is shorter than maxK or longer than maxK (Supplementary Table 1). We found no matter whether the DTR length is longer or shorter than the maxK, (1) if the genome did not contain another intra-genomic repeat longer than the maxK, all three assemblers will generate a single contig. (2) if the genome contains another intra-genomic repeat longer than the maxK, metaSPAdes will still generate a single contig, while IDBA_UD and MEGAHIT will obtain three contigs that broke at the intra-genomic repeat sites.

**Inter-genome shared regions.** We obtained two simulated genomes (genome 1 and 2) with a shared region longer than maxK in two cases, (1) coverage of genome 1 = 10 X, coverage of genome 2 = 30 X (Supplementary Fig. 3, upper panel), and (2) both genomes have a coverage of 20 X (Supplementary Fig. 3, bottom panel). In case (1), both IDBA_UD and MEGAHIT obtained five contigs, with the shared region represented by a single contig of 40 X coverage, and it could be used to concatenate the two contigs from genome 1 into a whole genome, and also the two contigs from genome 2 into a whole genome, and the contig end sequences are with an overlap of the expected length (maxK-1 for IDBA_UD and maxK for MEGAHIT), while metaSPAdes obtained three contigs, one of them represented the whole sequence of genome 2 (the dominant one), and the other two contigs represented the two parts of the genome 1 without the shared region. In case (2), IDBA_UD and MEGAHIT obtained the same contigs as in case (1), while metaSPAdes obtained the complete sequences of both genomes. These simulated results indicated that metaSPAdes is doing better than the other two assemblers in such cases.

We did not show here the simulated genomes with a shared region shorter than the maxK, as in this case, all three assemblers will obtain two complete genomes without fragmentation.

**Within-population variations.** For within-population variations, we obtained simulated genomes of two subpopulations (Supplementary Fig. 4) and three subpopulations (Supplementary Fig. 5).

For two subpopulations, if the variation region is shorter than maxK, a contig representing the more abundant subpopulation was generated by all three assemblers (Supplementary Fig. 4a). If the variation region is longer than maxK, IDBA_UD always breaks at the starting and ending points of the variable regions (Supplementary Fig. 4b). However, metaSPAdes and MEGAHIT generated different numbers of contigs that are dependent on the abundance of the subpopulations (reflected by the coverage of the simulated genomes), they generated fragmented contigs as

IDBA_UD did when the abundance of the two subpopulations was similar (for example, 50%/50%, 60%/40%), or generated a whole length of the more abundant subpopulation when the abundance of the two subpopulations was distinct enough (for example, 70%/30%, 80%/20%, 90%/10%). In some cases, metaSPAdes and MEGAHIT also generated the variation region as a contig of a less abundant subpopulation.

For three subpopulations, four different cases were tested with simulated genomes and reads. (1) In the first case, the starting and the ending sites of the variation region among the three subpopulations are the same (Supplementary Fig. 5a). No matter what the sequencing depths of the three subpopulations are, IDBA_UD always broke at the point of the variation starting and the ending sites, generating 5 contigs. While metaSPAdes obtained 5 contigs only when the sequencing depths were 50 X for all three subpopulations, in other assemblies, it obtained one full genome of the most dominant one, or three contigs (including the full genome of the most dominant one, each of the other two contigs represented the variable region of a less dominant subpopulation). For MEGAHIT, it obtained 5 contigs that broke at the point of the variation starting and the ending sites, in four of six of the tested sequencing depths combinations. In the other two sequencing depths combinations, MEGAHIT did the same as metaSPAdes, generating three contigs. (2) In the second case, the variation region starts at different sites and ends at the same site (which is the same end at different sites but starts at the same site, thus not shown here) (Supplementary Fig. 5b). (3) In the third and fourth cases (Supplementary Figs. 5c and d), the variation region starts at different sites and also ends at different sites. From the assemblies in these three cases, we could obviously conclude that, no matter what the sequencing depth combination is, IDBA_UD always broke at the variation points, generating the same number of contigs for all the assemblies in each of the three cases.

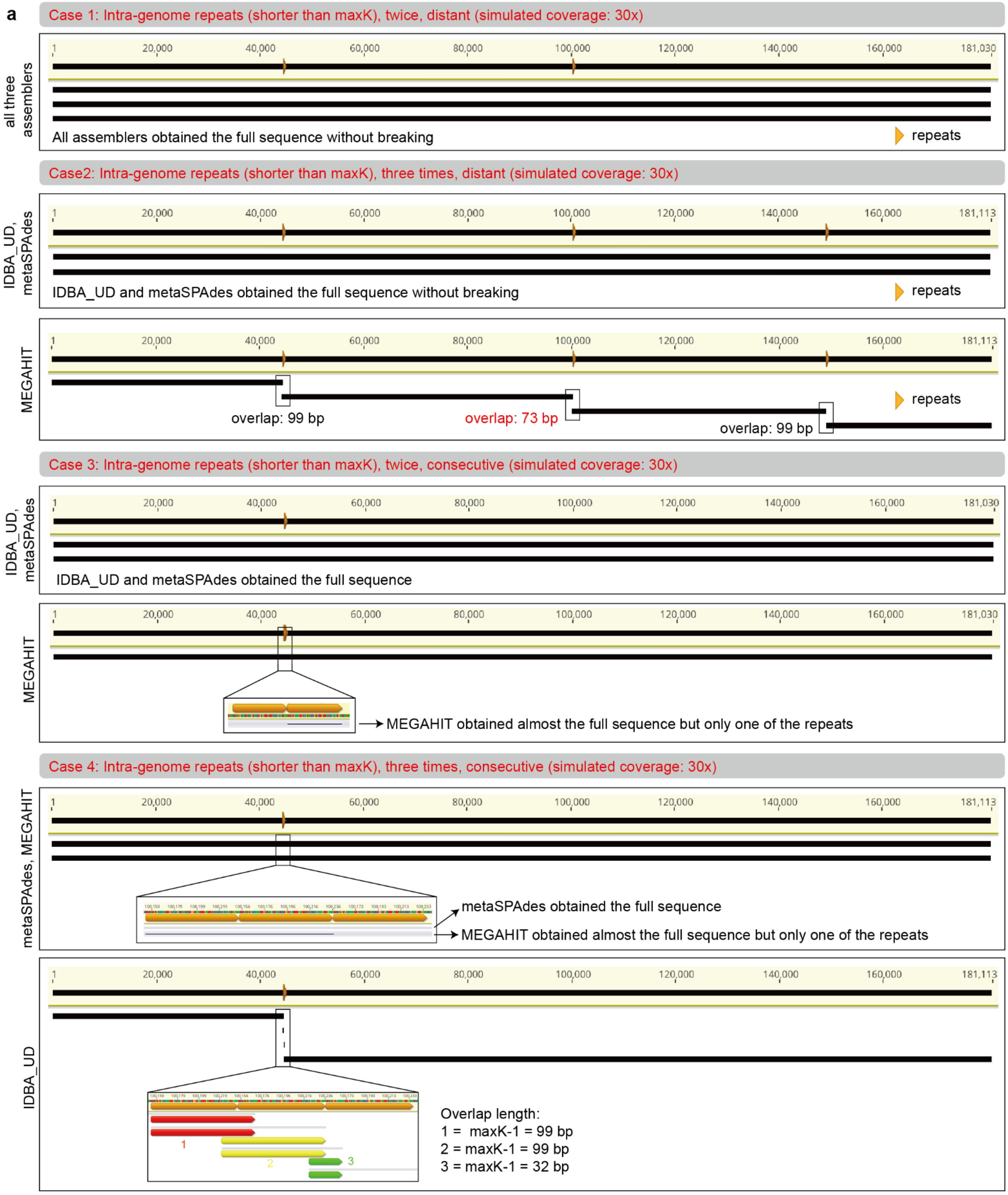


**Supplementary Fig. 1 | Simulated sequences with intra-genome repeats that are shorter than the largest kmer used in *de novo* assembly.** Different cases include: (case 1) repeats occurring twice and distant from each other, (case 2) repeats occurring triple times and distant from each other, (case 3) repeats occurring twice and consecutive with each other, and (case 4) repeats occurring triple times and consecutive with each other. The simulated genomes were in silico "sequenced" with a coverage of 30X, and the generated reads were assembled using metaSPAdes, IDBA_UD, and MEGAHIT in each case. If a given assembly obtained more than one contig, the overlapped sequence lengths (if detected) of the contigs are shown.

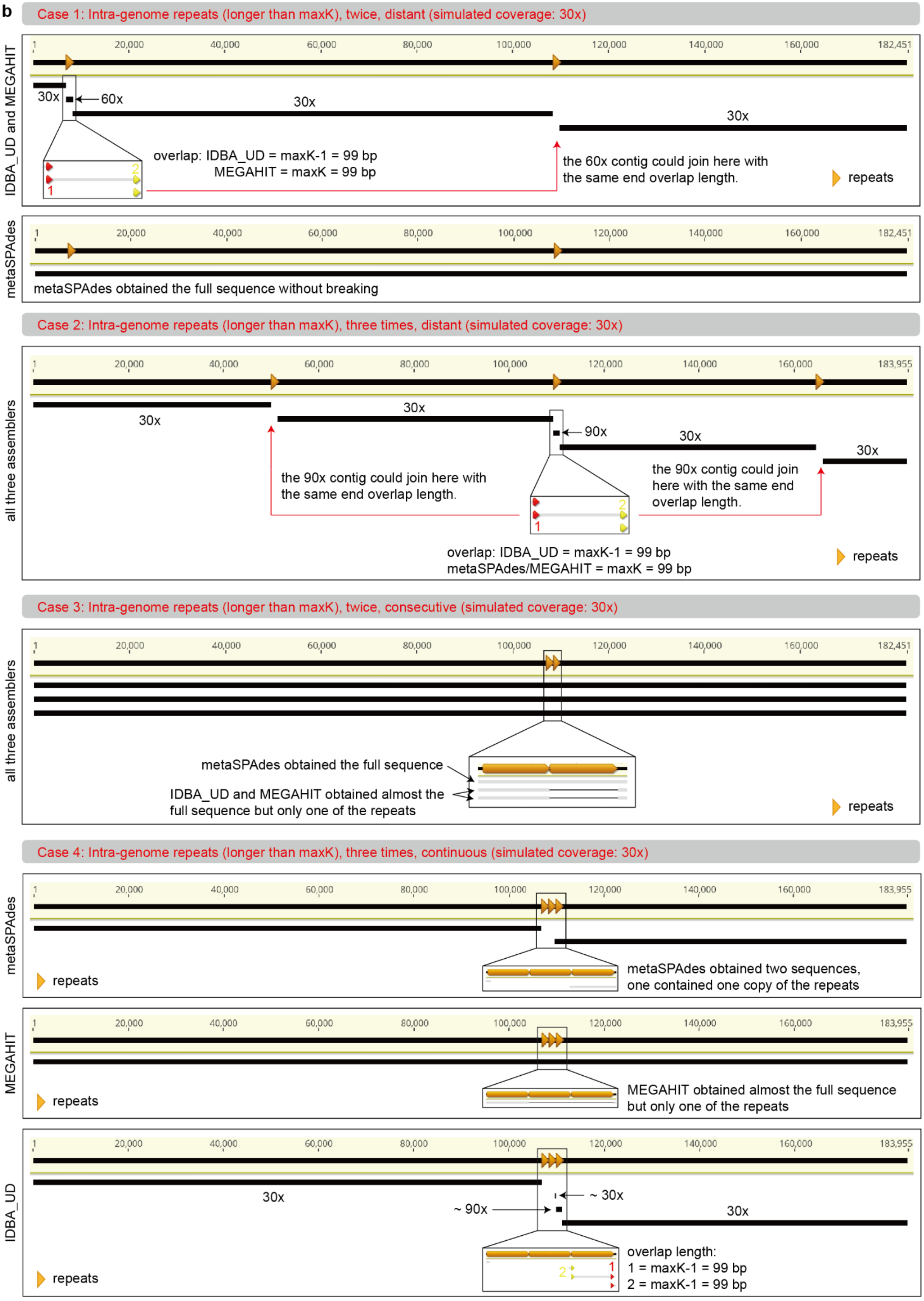


**Supplementary Fig. 2 | Simulated sequences with intra-genome repeats that are longer than the largest kmer used in *de novo* assembly.** Different cases include: (case 1) repeats occurring twice and distant from each other, (case 2) repeats occurring triple times and distant from each other, (case 3) repeats occurring twice and consecutive with each other, and (case 4) repeats occurring triple times and consecutive with each other. The simulated genomes were in silico "sequenced" with a coverage of 30X, and the generated reads were assembled using metaSPAdes, IDBA_UD, and MEGAHIT in each case. If a given assembly obtained more than one contig, the overlapped sequence lengths (if detected) of the contigs are shown.

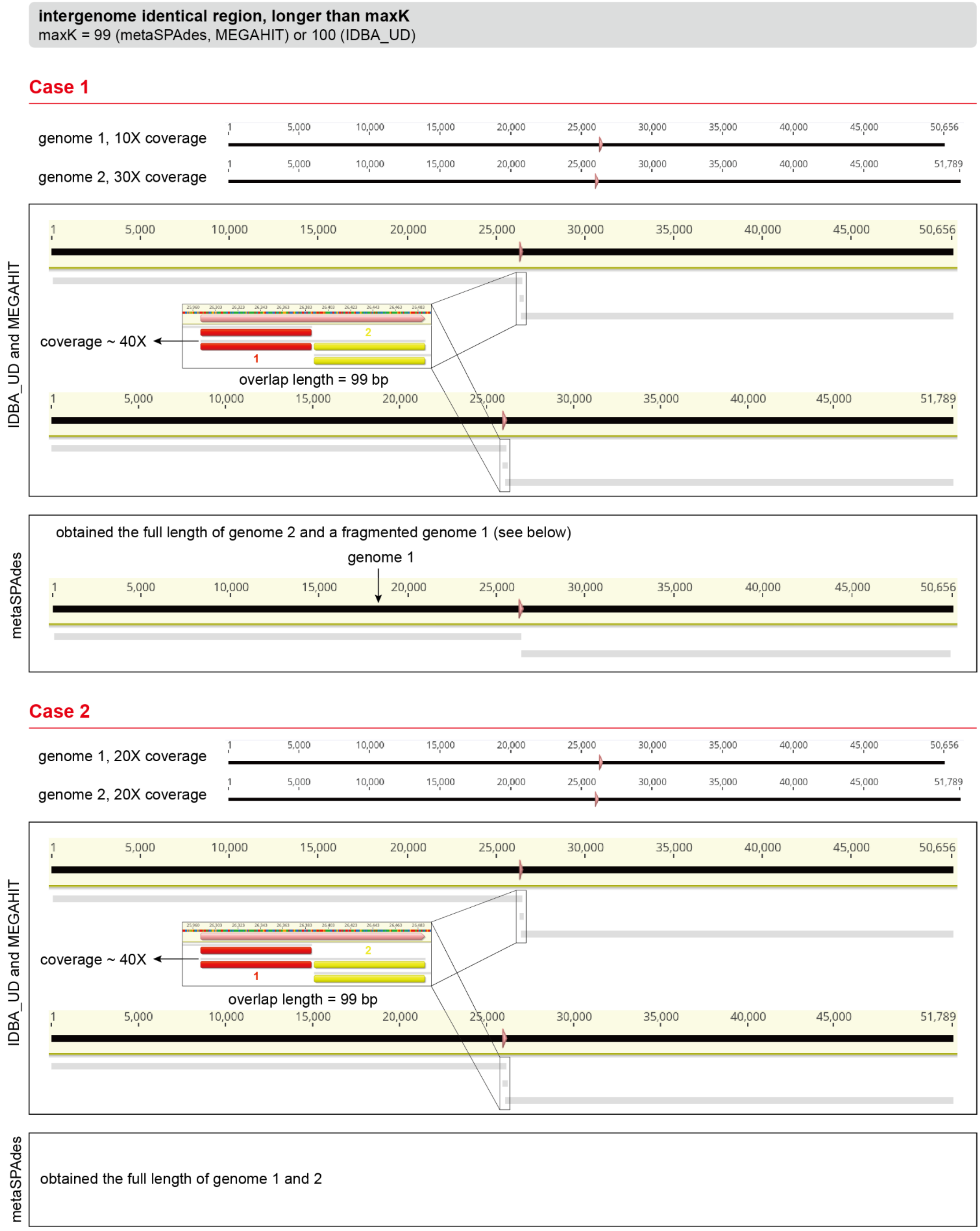


**Supplementary Fig. 3 | Simulated sequences with inter-genome shared regions.** Two different cases are shown: (case 1) where the two simulated genomes were in silico "sequenced" with coverage of 10X (genome 1) and 30X (genome 2), and (case 2) where both of the two simulated genomes were in silico "sequenced" with a coverage of 20X. The generated reads were assembled using metaSPAdes, IDBA_UD, and MEGAHIT in each coverage combination. If a given assembly obtained more than one contig, the overlapped sequence lengths (if detected) of the contigs are shown.

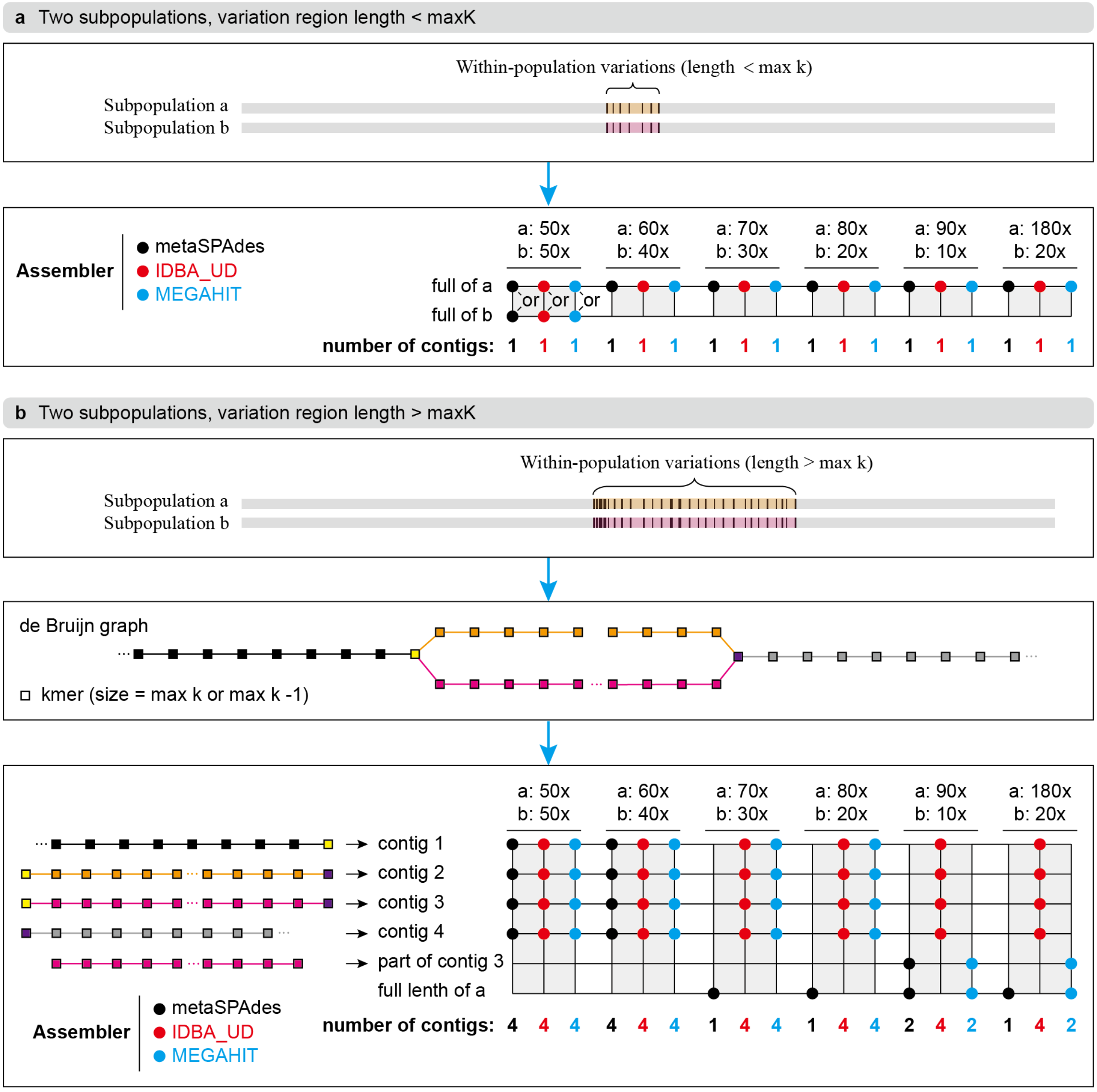


**Supplementary Fig. 4 | Simulated sequences with variations between two subpopulations.** The length of the variation region between the two subpopulations is compared for two cases: (a) when it is shorter than the maximum kmer (maxK) used in the assembly, and (b) when it is longer than the maxK used in the assembly. The simulated genomes were in silico "sequenced" with different coverage combinations, and the generated reads were assembled using metaSPAdes, IDBA_UD, and MEGAHIT in each coverage combination. The resulting contigs for each coverage combination are represented by colored circles.

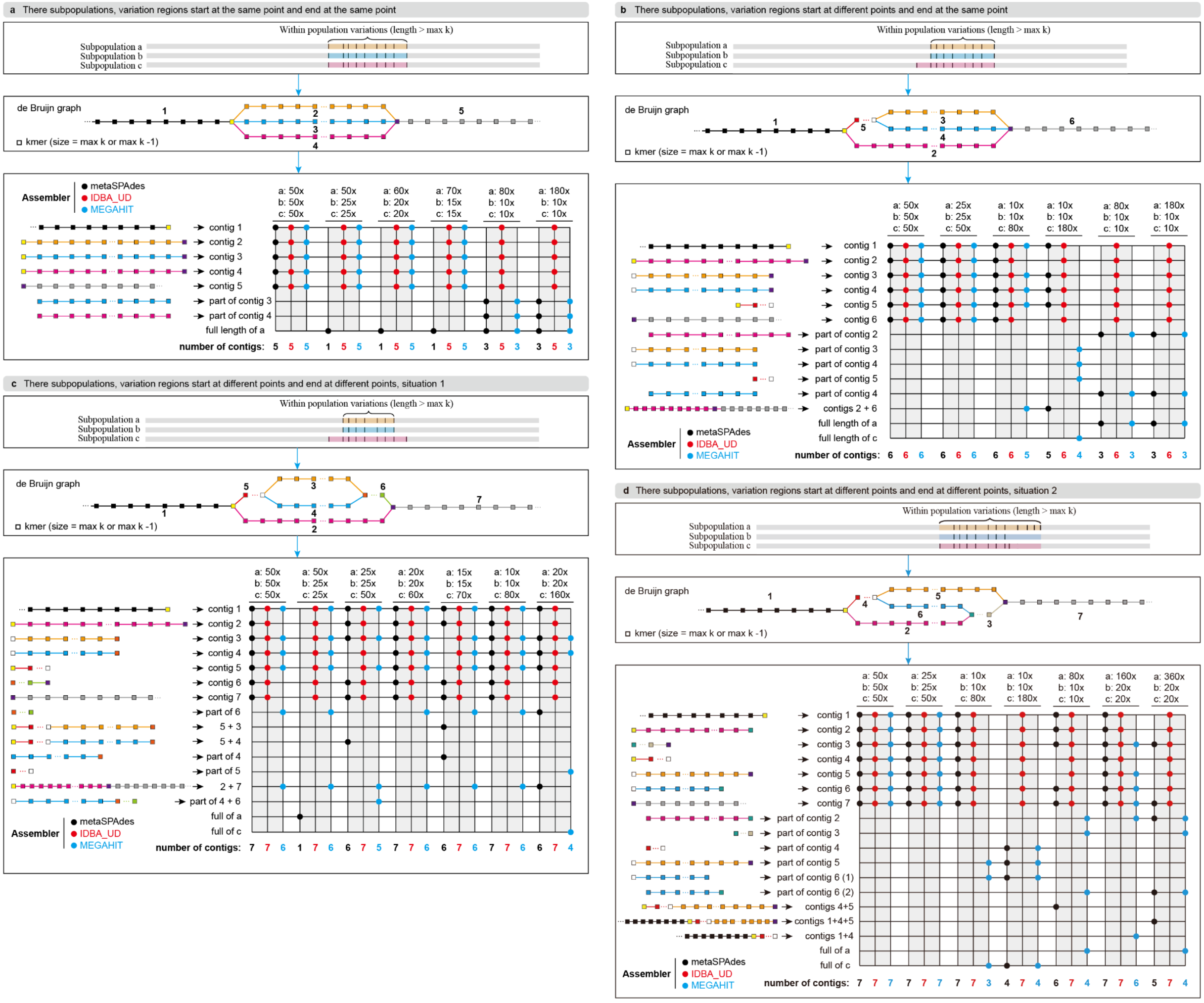


**Supplementary Fig. 5 | Simulated sequences with variations among three subpopulations.** The simulated genomes contain variation regions within three subpopulations, characterized by (a) identical start and end points, (b) different start points but the same end point, (c) different start and end points (situation 1), and (d) different start and end points (situation 2). The length of the variation regions exceeds the maximum kmer (maxK) used in the de novo assembly. The genomes were in silico "sequenced" with different coverage combinations, and the resulting reads were individually assembled using metaSPAdes, IDBA_UD, and MEGAHIT for each coverage combination. The contigs obtained from each coverage combination are represented by colored circles.

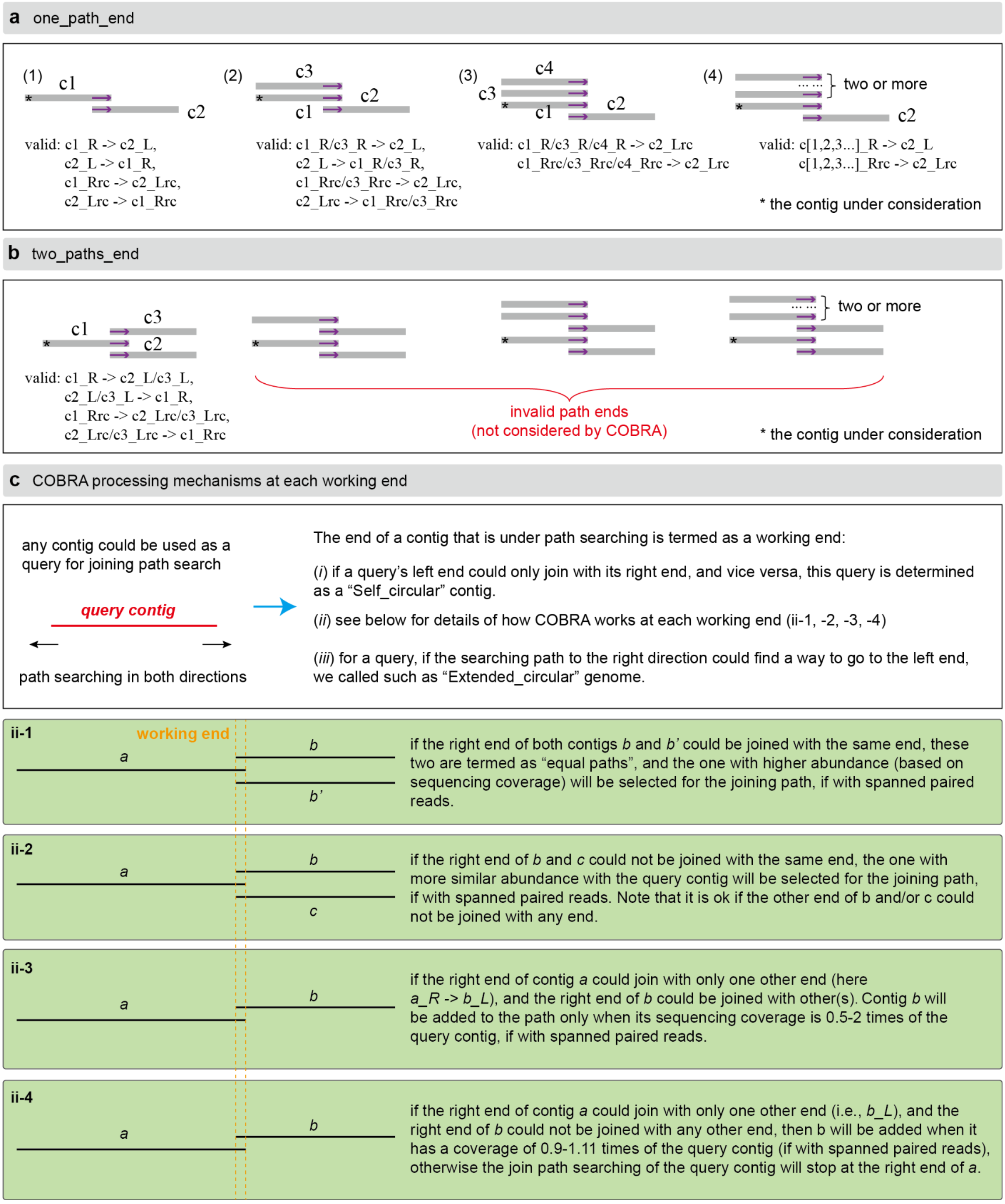


**Supplementary Fig. 6 | Detailed working principles of COBRA.** (a) and (b) illustrate the valid joins considered by COBRA during the joining path search for each query contig, specifically in the scenarios of "one_path_end" and "two_paths_end" respectively. In the diagrams, each gray bar represents a contig (e.g., c1 = contig_1), with its right sequence end labeled as "c1_R". The term "rc" denotes reverse complementary, and the arrow represents the contig end sequence, which is of length maxK or maxK-1. The subject contig under extension is marked with an asterisk on the contig bar, while all valid joins (direction sensitive) are displayed below the corresponding diagram. (c) A comprehensive overview of COBRA's working principles at each stage. In the case of "two_paths_end", the path with a closer coverage match to that of the "query contig" will be selected.

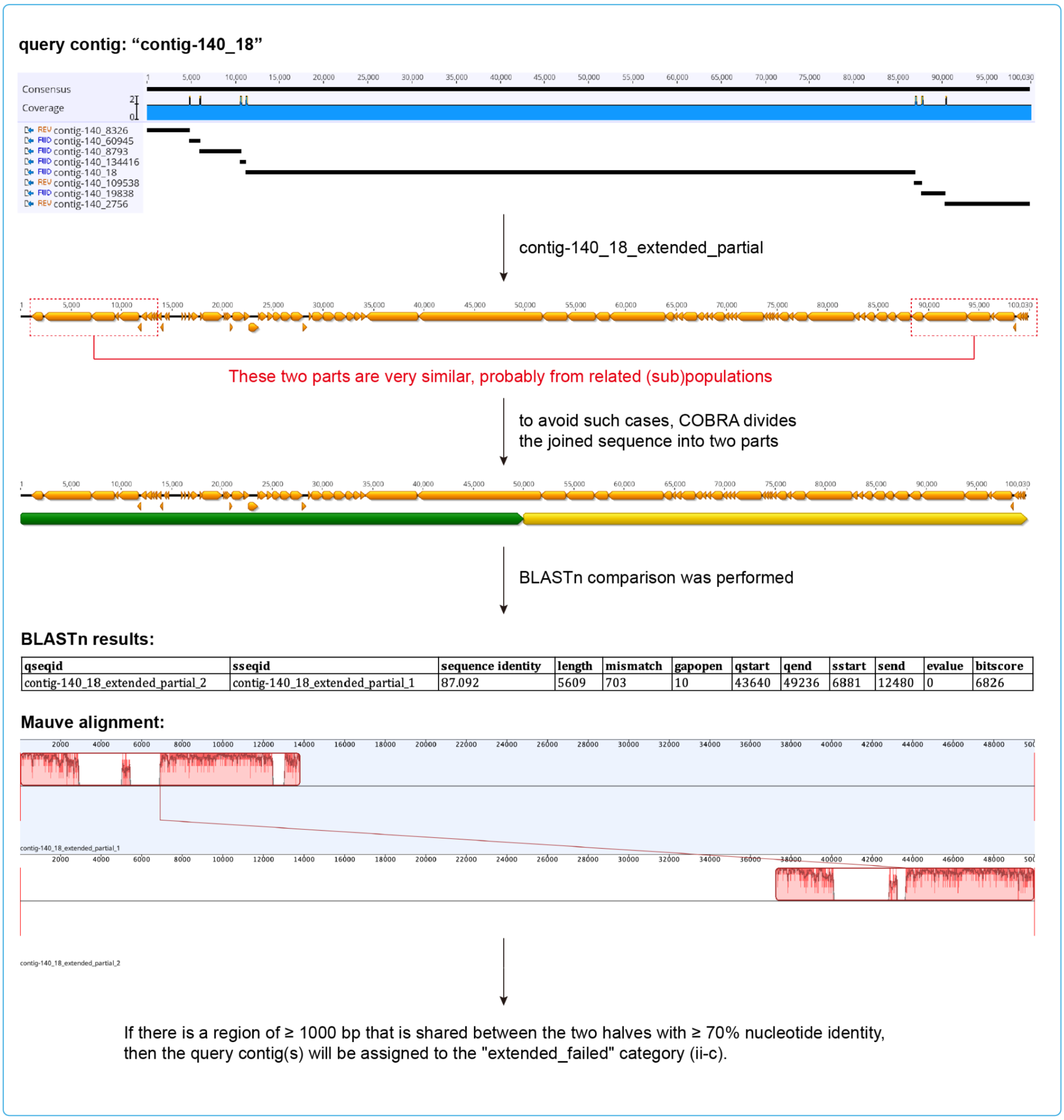


**Supplementary Fig. 7 | Screening of contigs joined from closely related genomes using BLASTn comparison.** To prevent the joining of fragmented contigs from closely related (sub)populations, a BLASTn comparison is conducted between the first half and second half of each joined COBRA sequence. If the two parts share a region with a minimum length of 1000 bp and a minimum nucleotide similarity of 70%, all the query contigs involved in the join are labeled as "extended_failed" to indicate the failed extension.

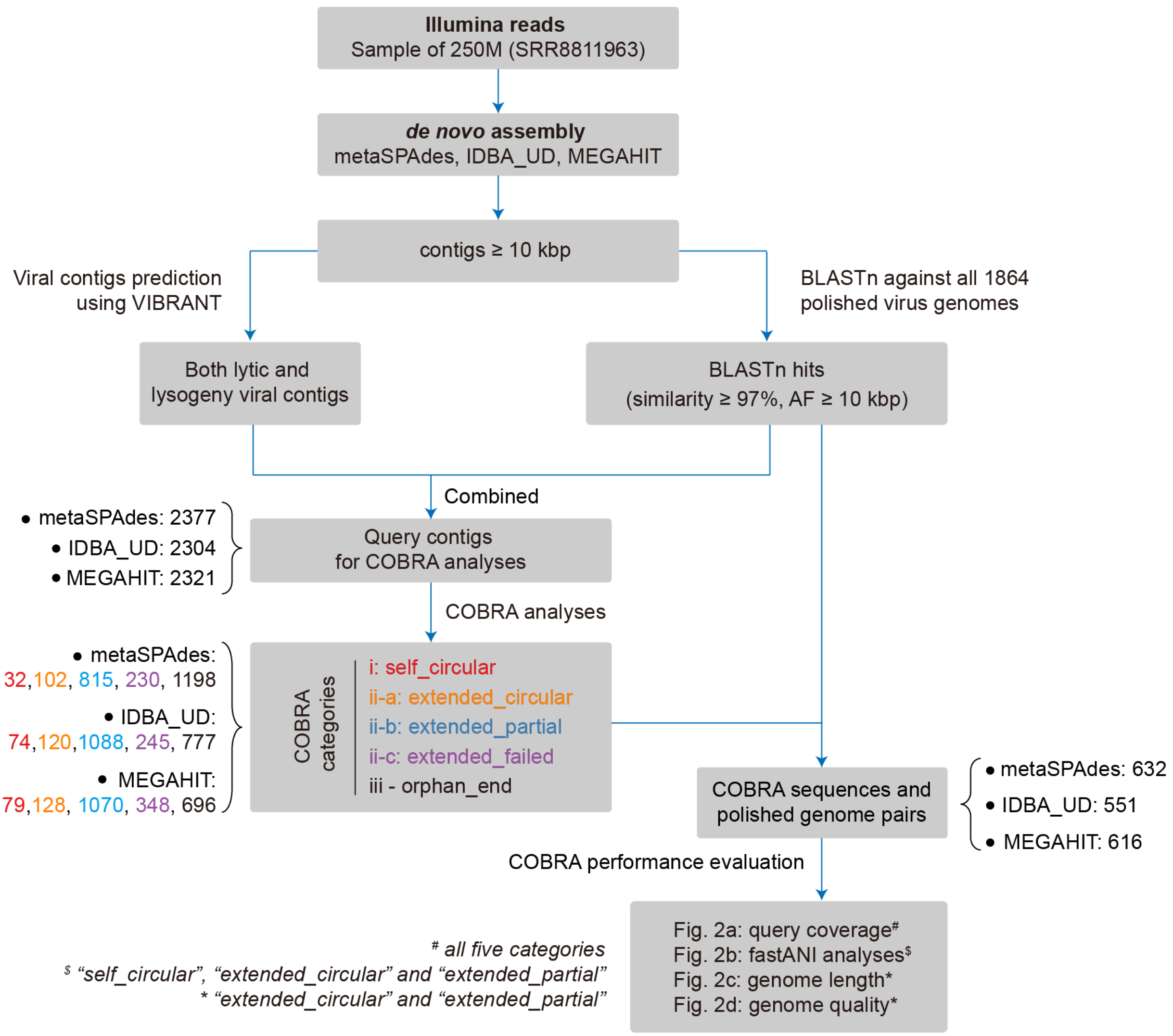


**Supplementary Fig. 8 | The pipeline to evaluate the performance of COBRA using a published virome sample.** The dataset was previously published by Beaulaurier et al. (2019). The raw Illumina reads were downloaded from NCBI, and quality control was conducted (see **Methods** in the main text). The quality paired-end reads were respectively assembled using metaSPAdes, IDBA_UD and MEGAHIT. The assembled contigs with a minimum length of 10 kbp were extracted for viral prediction (with VIBRANT) and searched against the polished viral genomes (see Beaulaurier et al. 2019 for details) for viral contigs (with BLASTn, nucleotide similarity ≥ 97%, aligned fraction ≥ 10 kbp). The obtained viral contigs were combined as queries for COBRA analyses. The "extended_circular" or "extended_partial" COBRA sequences were compared with the corresponding polished genomes (determined by the abovementioned BLASTn search) to evaluate the performance of COBRA (see results in **Fig. 2** of the main text).

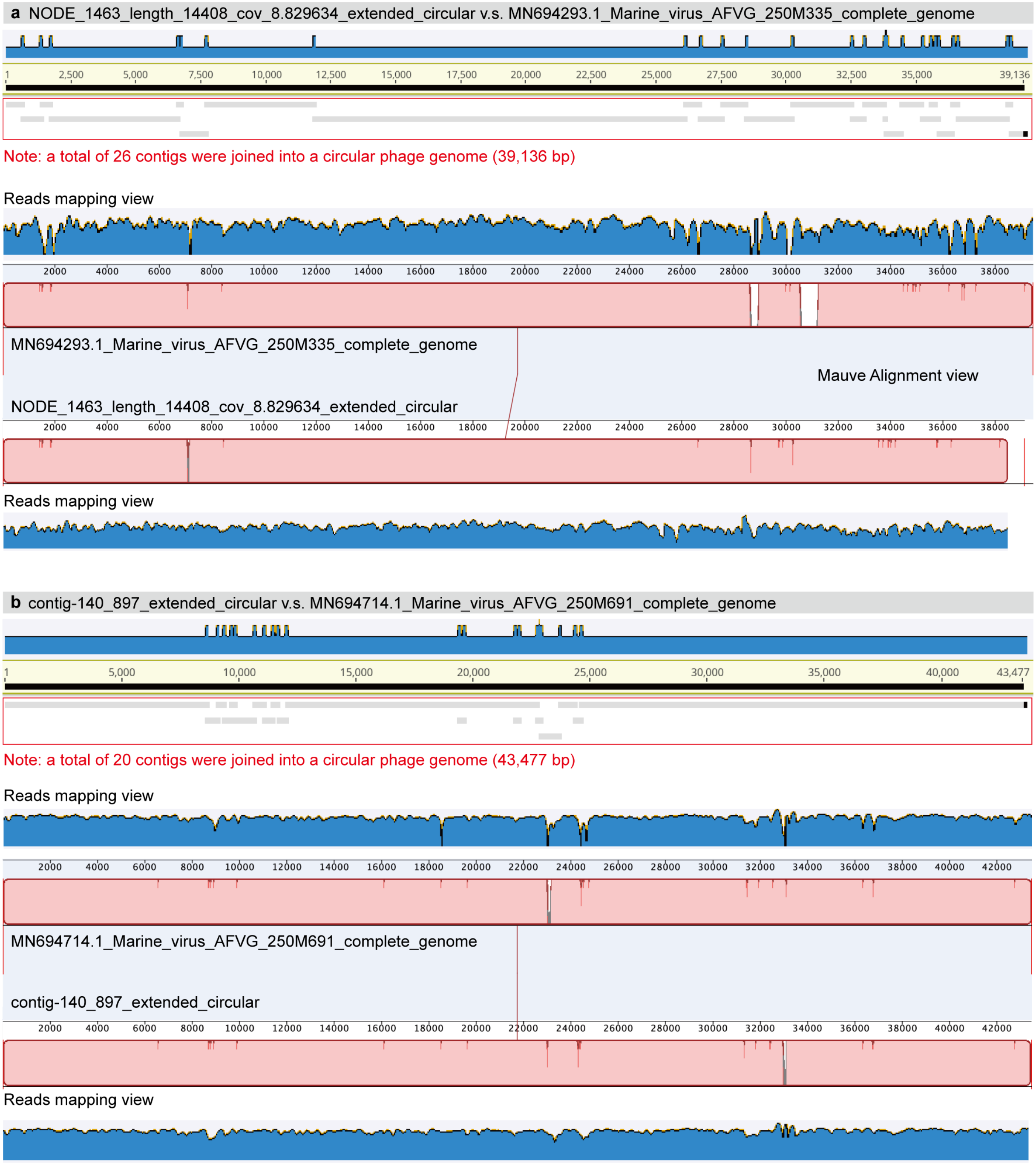


**Supplementary Fig. 9 | Circular extension of query virus contigs by COBRA.** Illustrative examples of genomes joined by COBRA that exhibit similarity to a polished genome, with contigs assembled using (a) metaSPAdes and (b) IDBA_UD. The Mauve Alignment view showcases the comparison between the COBRA “extended_circular” genome and the corresponding polished genome, along with their respective read mapping profiles for validation.

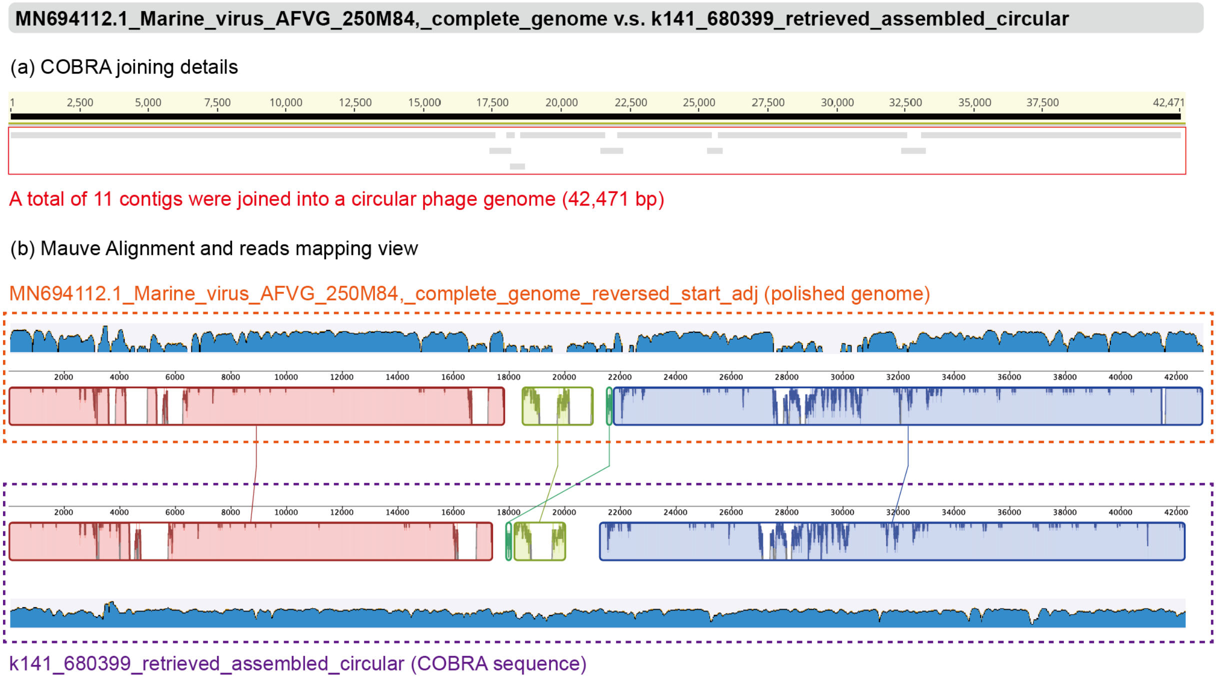


**Supplementary Fig. 10 | One “extended_circular” from the MEGAHIT assembly with < 90% “AF_COBRA” and < 90% “AF_polished”.** (a) The original contig was assembled by MEGAHIT, which was extended to a circular genome including a total of 11 contigs. (b) The mauve alignment and reads mapping profiles show that the COBRA genome represents the dominant subpopulation while the polished genome represents the rare subpopulation in the sample.

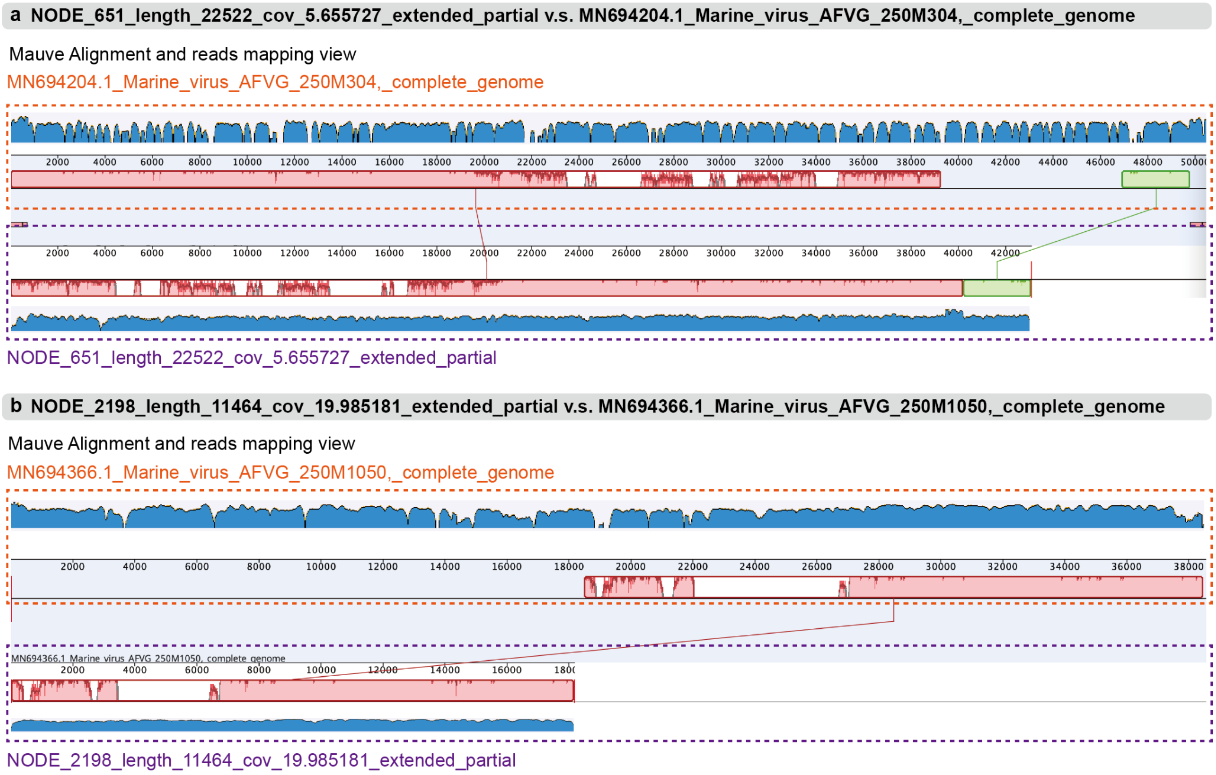


**Supplementary Fig. 11 | Examples of COBRA and polished genome pairs with relatively lower "AF_COBRA" values.** To document that such pairs are due to variations between subpopulations, the mauve alignment and read mapping profiles for each pair of the polished genome and COBRA sequence are shown.

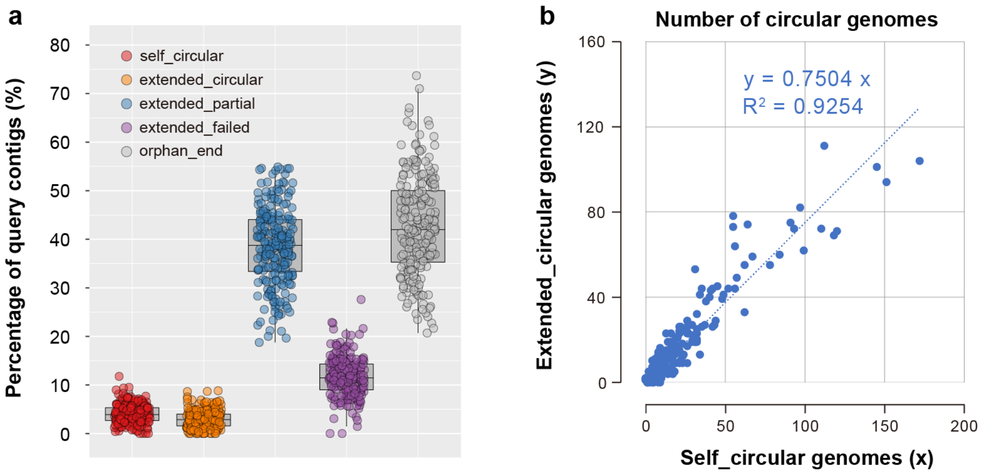


**Supplementary Fig. 12 | COBRA analyses of 231 freshwater metagenomes. (a)** The percentage of queries in each COBRA category. **(b)** The comparison of the number of "self_circular" and unique "extended_circular" genomes from freshwater metagenomes. Interestingly, the number of "self_circular" genomes is highly related to that of unique "extended_circular" genomes.

##### (a) Zhao et al., 2022, HugePhage 1, 852 kbp

- evaluated by viral identification tools

CheckV: viral_genes = 33, host_genes = 533, provirus = yes, termini = complete-prophage, viral_length = 67,045 bp, viral region = 487042-554086

VIBRANT: viral length = 84,279 bp, life style = lysogenic, viral region = ~490936 - ~584027 bp

- alignment with contigs from Delftia acidovorans strain 2167 DR66.Contig240 (a bacterial genome; NCBI ID = JOUB01000003)

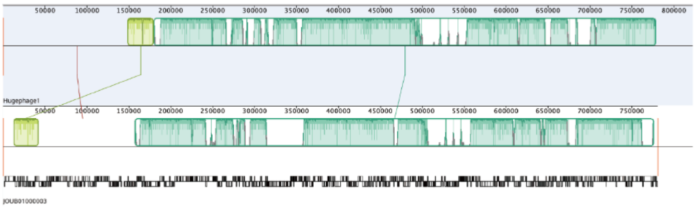


Based on all the information we obtained here, the HugePhage 1 sequence contains a prophage genome (67-84 kbp in length), given the genome wide alignment with the contigs of a Delftia acidovorans genome from NCBI, and the high similarity of large terminase protein with those from Delftia spp., we conclude that "HugePhage 1" should be a small phage of Delftia member.

- high similarity of HugePhage 1 large terminase to those of Delftia spp.

| | Description | Scientific Name | Max Score | Total Score | Query Cover | E value | Per. Ident | Acc. Len | Accession |
|---|---|---|---|---|---|---|---|---|---|
| ☑ | terminase family protein [Delftia sp. ZNC0008] | Delftia sp. ZNC0008 | 1040 | 1040 | 100% | 0.0 | 99.42% | 515 | WP_047469065.1 |
| ☑ | terminase family protein [Delftia sp. RIT313] | Delftia sp. RIT313 | 1040 | 1040 | 100% | 0.0 | 99.42% | 515 | WP_043821036.1 |
| ☑ | terminase family protein [Delftia sp. ZNC0008] | Delftia sp. ZNC0008 | 1035 | 1035 | 100% | 0.0 | 98.64% | 515 | WP_047473737.1 |
| ☑ | terminase family protein [Delftia tsuruhatensis] | Delftia tsuruhatensis | 1034 | 1034 | 100% | 0.0 | 98.45% | 515 | WP_046984607.1 |
| ☑ | terminase family protein [Delftia acidovorans] | Delftia acidovorans | 1033 | 1033 | 100% | 0.0 | 98.45% | 515 | WP_034395113.1 |
| ☑ | terminase family protein [Delftia acidovorans] | Delftia acidovorans | 1033 | 1033 | 100% | 0.0 | 98.45% | 515 | WP_097203122.1 |
| ☑ | terminase [Delftia acidovorans] | Delftia acidovorans | 1032 | 1032 | 100% | 0.0 | 98.45% | 515 | WP_239971178.1 |
| ☑ | terminase family protein [Delftia tsuruhatensis] | Delftia tsuruhatensis | 1032 | 1032 | 100% | 0.0 | 98.45% | 515 | WP_133073502.1 |
| ☑ | terminase [Delftia acidovorans] | Delftia acidovorans | 1032 | 1032 | 100% | 0.0 | 98.25% | 515 | PJO37630.1 |
| ☑ | terminase family protein [Delftia tsuruhatensis] | Delftia tsuruhatensis | 1032 | 1032 | 100% | 0.0 | 98.25% | 515 | WP_046239682.1 |
| ☑ | terminase family protein [Delftia acidovorans] | Delftia acidovorans | 1030 | 1030 | 100% | 0.0 | 98.25% | 515 | WP_180319372.1 |
| ☑ | terminase family protein [Delftia acidovorans] | Delftia acidovorans | 1029 | 1029 | 100% | 0.0 | 98.06% | 515 | MBN9323155.1 |
| ☑ | terminase family protein [Delftia acidovorans] | Delftia acidovorans | 1028 | 1028 | 100% | 0.0 | 98.06% | 515 | WP_197956440.1 |
| ☑ | terminase family protein [Delftia sp. SD018] | Delftia sp. SD018 | 1028 | 1028 | 100% | 0.0 | 98.06% | 515 | WP_207475865.1 |

##### (b) Zhao et al., 2022, HugePhage 2, 839 kbp

- evaluated by viral identification tools

CheckV: viral_genes = 37, host_genes = 533, provirus = yes, termini = complete-prophage, viral_length = 27,719 bp, viral region = 154998-182716

VIBRANT: viral length = 32,493 bp, life style = lysogenic, viral region = ~148362 - ~183490 bp

- alignment with contigs from Ruminococcus sp. CAG:108 (a bacterial genome; NCBI ID = CAXD010000000)

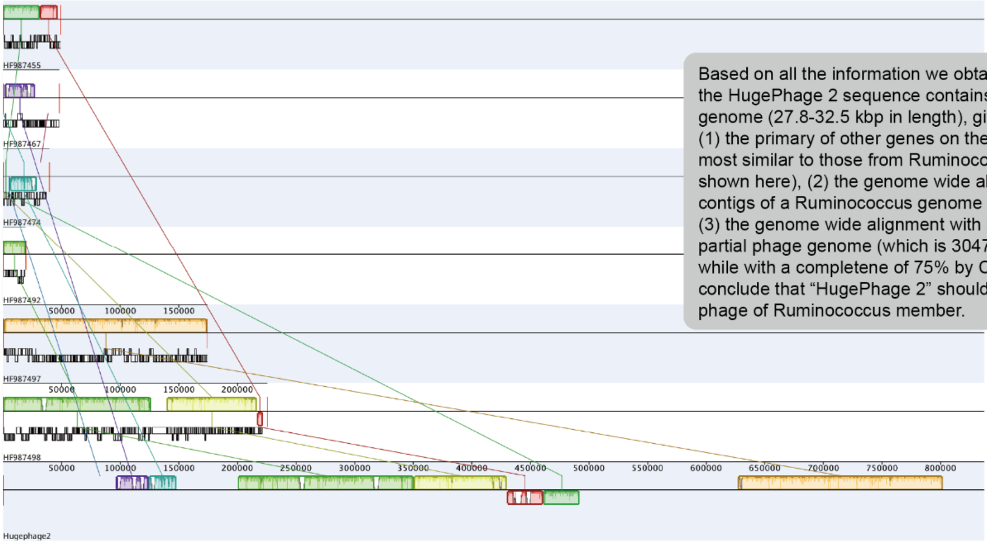


Based on all the information we obtained here, the HugePhage 2 sequence contains a prophage genome (27.8-32.5 kbp in length), given that (1) the primary of other genes on the sequence were most similar to those from Ruminococcus sp. (not shown here), (2) the genome wide alignment with the contigs of a Ruminococcus genome from NCBI, and (3) the genome wide alignment with a published partial phage genome (which is 30475 bp in length, while with a completene of 75% by CheckV), we conclude that "HugePhage 2" should be a small phage of Ruminococcus member.

- alignment with the partial genome of Caudoviricetes sp. isolate ctxHS4 (NCBI ID = BK019798), whose host is Ruminococcus (Tisza and Buck, 2021)

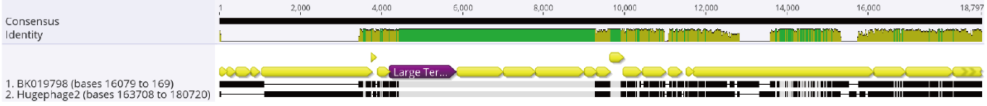


see the text legend on the next page.

**Supplementary Fig. 13 | Evidence indicating that the two published huge phages with genome sizes over 800 kbp were prophage sequences.** (a) HugePhage 1 and (b) HugePhage 2 reported by Zhao et al. (Zhao et al. 2022). The genomes were downloaded via the link provided in the corresponding literature, and evaluated using the viral identification tools of CheckV (Nayfach et al. 2020) and VIBRANT (Kieft et al. 2020) to obtain the related information, including “viral_length”, “viral region”, “life style” and others. The genomes were searched against NCBI for highly similar sequences, which led to the identification of the corresponding bacterial genomes (and one phage genome for HugePhage 2) as shown in the figure. The protein-coding genes predicted using Prodigal (-m -p single) (Hyatt et al. 2010) from the two phage genomes were searched against the RefSeq database (Pruitt et al. 2007) using BLASTp for their taxonomic assignment. According to the corresponding literature of Caudoviricetes sp. isolate ctxHS4 (NCBI ID = BK019798), i.e., Tisza and Buck (Tisza and Buck 2021), the bacterial host of this phage is predicted as “Ruminococcus”.

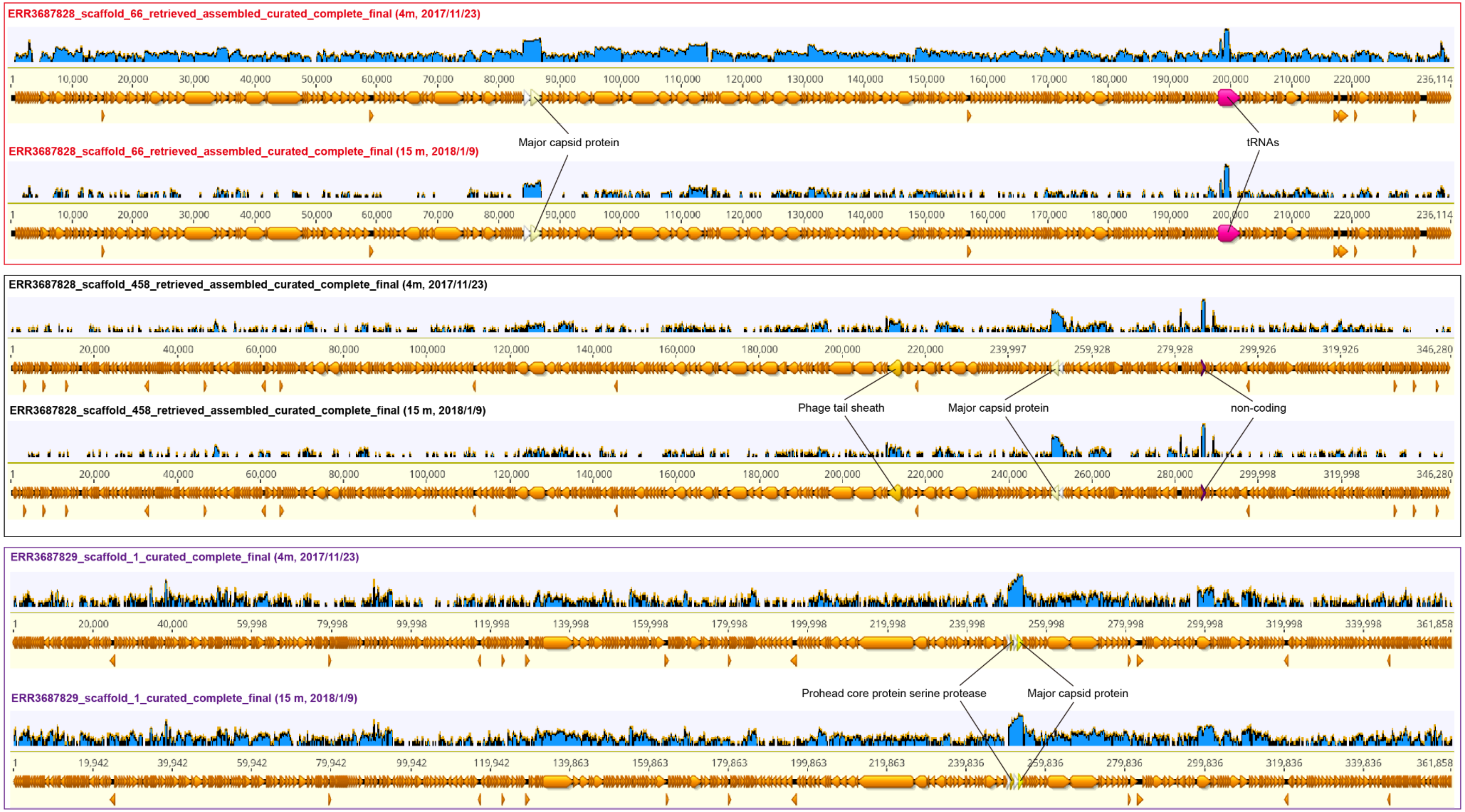


**Supplementary Fig. 14 | Examples of the in situ gene expression of huge phages with genomes reconstructed from Lake Rotsee.** The reads from different metatranscriptomic samples were mapped to each of the three genomes using Bowtie2 with default parameters. Then the bam files were imported to Geneious and remapped allowing no mismatch for each read. The huge phage genome names, sampling depths, and sampling time points are shown. The most highly transcribed ones are highlighted with their annotations.

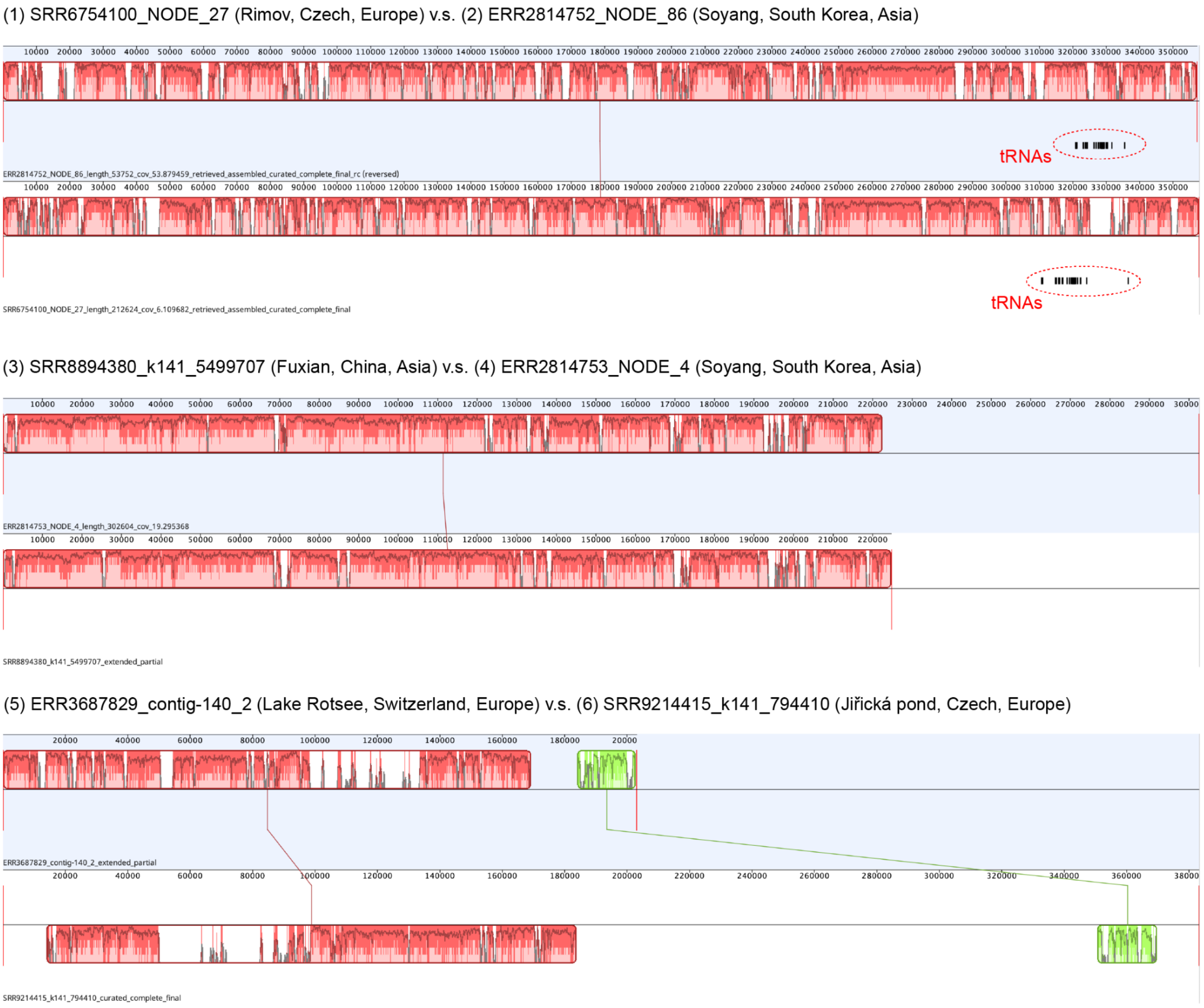


**Supplementary Fig. 15 | Mauve alignment of similar huge phage genomes from distant sampling sites.** The alignments of huge phage genomes were reconstructed from freshwater lakes of different countries. The tRNAs in the first pair, i.e., genome (1) and (2) were highlighted with dashed circles.

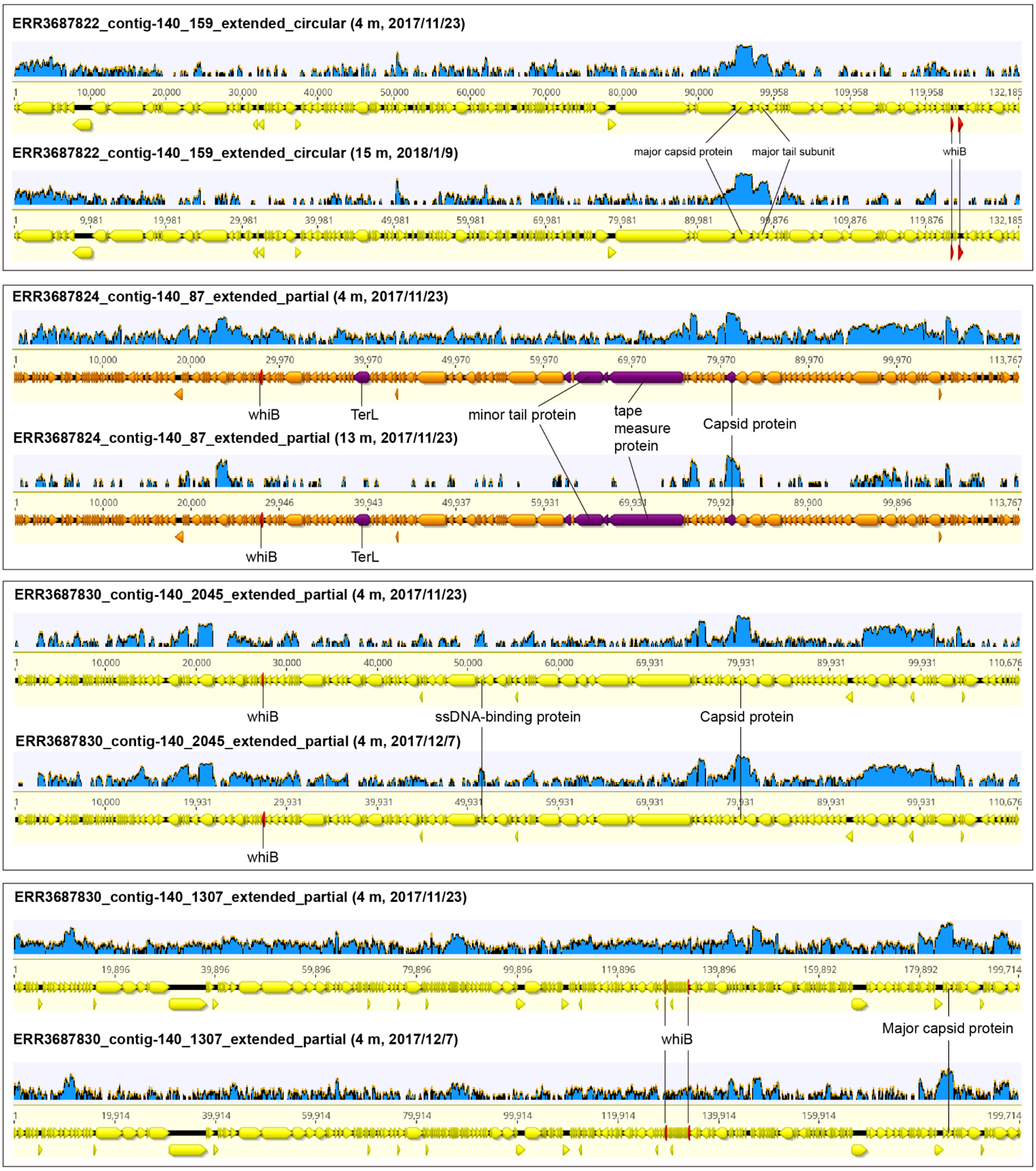


**Supplementary Fig. 16 | Examples of the *in situ* gene expression of whiB-encoding actinophages with genomes reconstructed from Lake Rotsee.** The reads from different metatranscriptomic samples were mapped to each of the four genomes using Bowtie2 with default parameters. Then the bam files were imported into Geneious and remapped allowing no mismatch for each read. The whiB genes are highlighted in red, some core phage structural protein genes are indicated with their annotations shown. The genome names, the sampling depths, and the sampling dates of the corresponding transcriptomic sample are shown at the top of each profile. A total of four actinophage genomes are shown (the one with two whiB genes is shown at the bottom), with each profiled with two transcriptomic samples.

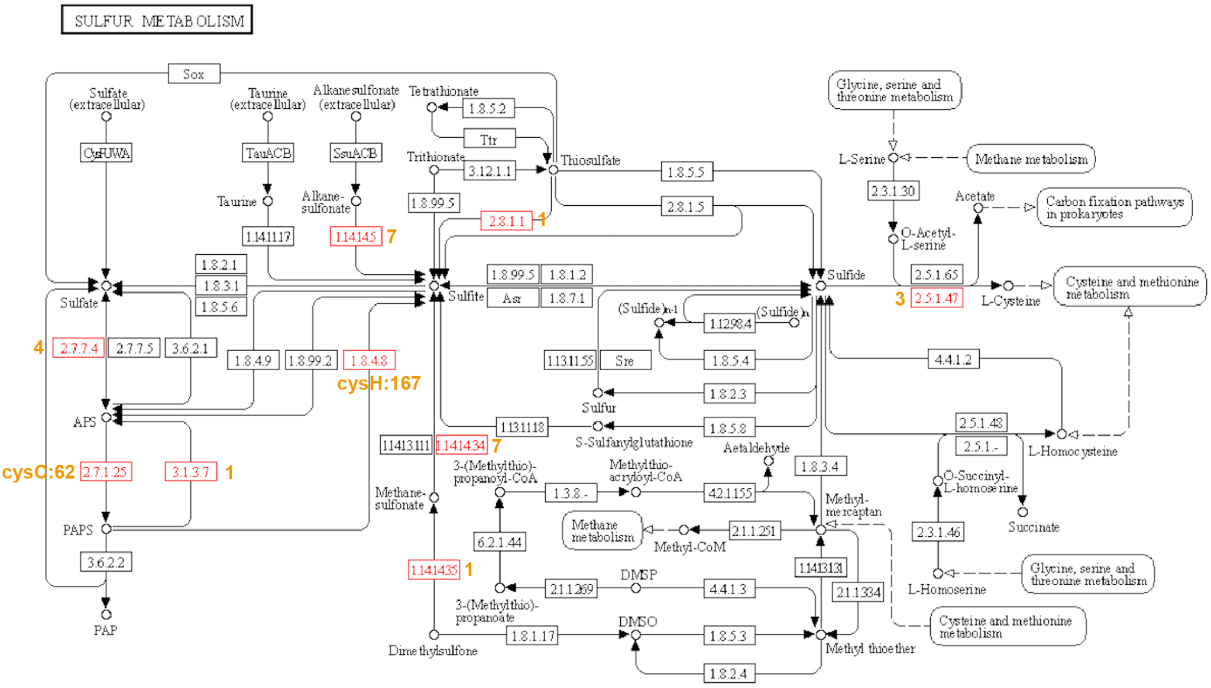


**Supplementary Fig. 17 | The number of sulfur metabolism-related AMGs detected in the phage genomes.** The detected genes are highlighted in red, and the corresponding number of genes is shown.

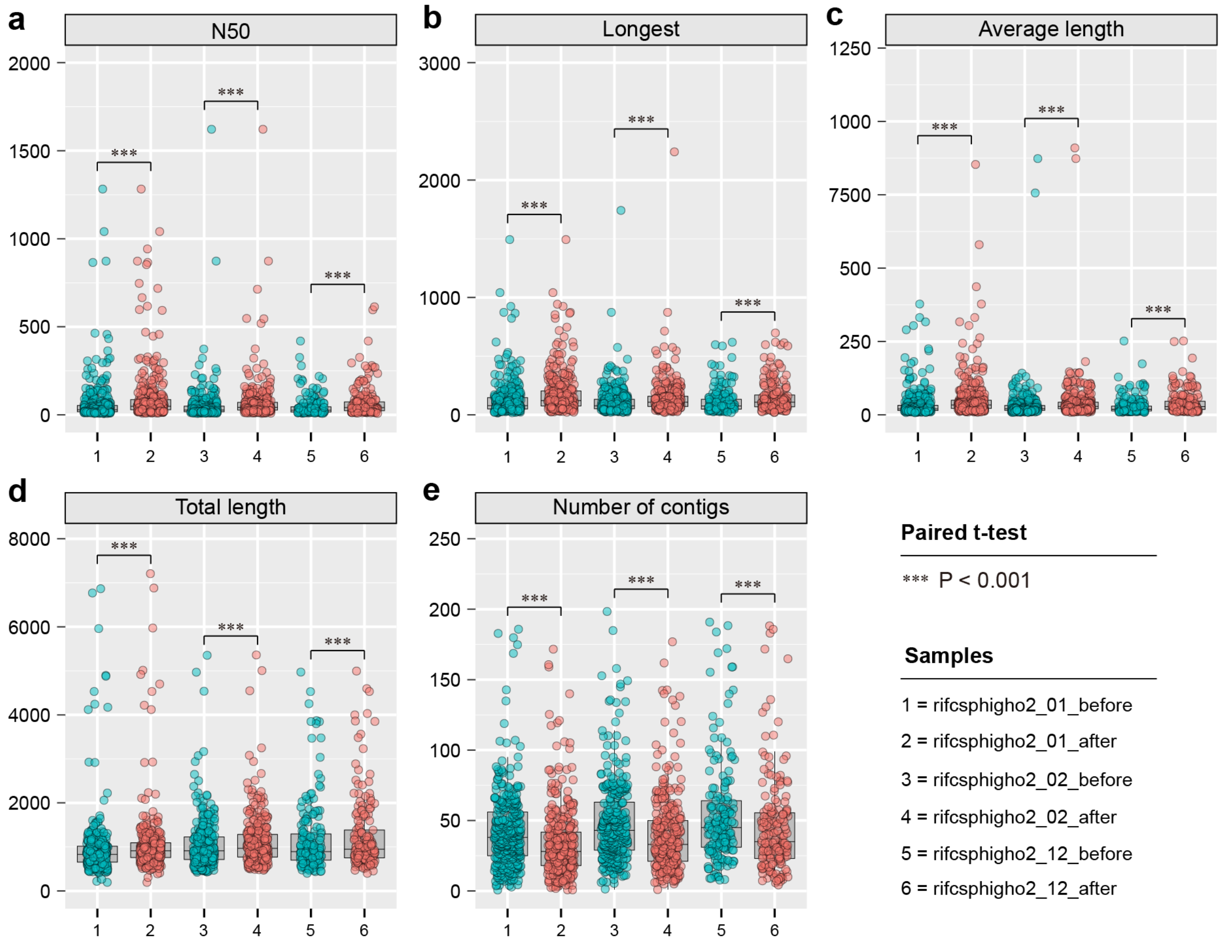


**Supplementary Fig. 18 | The performance of COBRA on non-viral genomes.** Comparison of MAGs before and after COBRA analyses regarding (a) N50, (b) largest length (i.e., longest), (c) average length, (d) total length, and (e) number of contigs. All the numbers are shown in kbps. For the samples, "_before" means the MAGs before the COBRA analyses (original bins), and "_after" means the MAGs post the COBRA analyses.
